# The specific roles of renal macrophages in monitoring and clearing off intratubular particles

**DOI:** 10.1101/2022.04.11.487834

**Authors:** Jian He, Yangyang Cao, Qian Zhu, Xinge Wang, Guo Cheng, Qiang Wang, Fei Han, Peng Shi, Xiao Z Shen

**Affiliations:** Department of Physiology and Department of Cardiology of the Second Affiliated Hospital, Zhejiang University School of Medicine, Hangzhou, Zhejiang, China; Department of Cardiology of the Second Affiliated Hospital, and Institute of Translational Medicine, Zhejiang University School of Medicine, Hangzhou, Zhejiang, China; Department of Laboratory Medicine, Zhejiang Hospital, Zhejiang University School of Medicine, Hangzhou, Zhejiang, China; Kidney Disease Center, the First Affiliated Hospital, Zhejiang University School of Medicine, Hangzhou, Zhejiang, China

## Abstract

During the filtrate of the glomerulus flows though the renal tubular system, a variety of microscopic sediment particles, including mineral crystals resulting from urine concentration, are generated. Dislodging these particles in the intratubular compartment is critical to ensure free flow of filtrate and the final formation of urine. However, the underlying mechanism for the clearance is unclear. Here, using high-resolution microscopy, we uncovered that the juxtatubular macrophages in the medulla constitutively formed transepithelial protrusions and were “sampling” urine contents. These behaviors were strengthened in the development of nephrolithiasis. In particular, the juxtatubular macrophages were efficient in sequestering and phagocytosing intraluminal sediment particles, and occasionally making transmigration to the tubule lumen to escort the excretion of urine particles. Specific depletion of renal macrophages precipitated kidney stone formation and aggravated the accompanied inflammation upon hyperoxaluria challenge. Thus, renal macrophages undertake a specific role in maintaining the tubular system unobstructed.

## INTRODUCTION

The kidney tubular system displays a hierarchically organized architecture which extends from the corpuscle, the initial filtering component exclusively distributing in the cortex, and collects and drains urine to the collecting duct, and eventually empties into the renal pelvis ^[1]^. The renal tubule is made up of a single layer of epithelial cells resting on a basement membrane. During urine passage through distinctive segments of renal tubule, dynamic reabsorption and excretion are executed by tubular epithelial cells ^[2]^. From cortex to medulla, there is a gradual osmolarity increase with the highest osmolarity at its deepest medulla, which is critical for urine concentration ^[3, 4]^. The complex organization of the kidney tissue is further reflected by the regional variations of the tubular epithelial cells as different segments express distinguishable transporters specified for reabsorption or secretion of certain materials ^[1, 5]^.

Although the physicochemical property of filtration barrier in the corpuscle precludes the passage of large blood-borne particles (e.g., blood cells) into the urine, the urine typically contains a significant amount of aggregated particles derived from varying sources. For instance, during the passage of renal tubule, about 99%of water in the corpuscle filtrate is reabsorbed so that the urine is significantly concentrated ^[1]^, which occasionally leads to supersaturation of some solutes in the distal nephron ^[5]^. The levels of urinary supersaturation correlate with nephrolithiasis (formation of kidney stone), and up to a quarter of fresh urine samples from healthy subjects show overt crystalluria ^[6]^. Second, apoptotic tubular epithelial cells routinely slough into the lumen of the renal tubules. It was estimated that approximately 70,000 cells of tubular origin are excreted in the urine per hour in human kidney ^[7]^. In addition, decrease in urine flow, increased acidity, and/or the presence of abnormal ionic or protein constituents expedites sedimentary particle formation in the distal segments of the renal tubule system ^[8–10]^. These small particles agglomerate together to form larger clusters. In agreement, the distal convoluted tubule and collecting duct are the principal sites of cast formation ^[11]^. Although millions of microscopic particles and large casts may form within the kidneys daily ^[12]^, most are excreted safely in the urine. However, diet and numerous pathologies greatly influence the range of crystal and cast burden that the kidneys experience ^[13, 14]^. Although the etiology of nephrolith and the formation of casts have been extensively studied ^[14–17]^, Whether there is an inherent “dislodging” mechanism, in addition to the urine flush, at paly for the removal of sedimentary particles is unclear.

Among the immune cells, macrophages (MØ) are the most frequent cell type within the barrier tissues. MØ are adapted to microanatomical structures, and these fundamental features are particularly pertinent for barrier epithelia of the skin, airway, gut, and genital tract that form the interface between the environment and the body, where they play roles more than immune surveillance ^[18–21]^. How epithelial barrier and macrophages are organized differs depending upon the tissue, its principal functions, and the specific stresses it endures. It was generally thought that the renal tubular epithelium generates a shield of sealed barrier to insulate the kidney from biotic and abiotic stresses derived from the urine ^[22–24]^, and phagocytes distribute underneath the epithelial layer in steady state ^[25, 26]^. In this study, we uncover that kidney medullary MØ constitutively extend transcellular protrusions through tubular epithelial monolayer, whereby MØ were sampling and monitoring urine contents. More importantly, these innate behaviors of MØ were crucial for keeping tubules unobstructed by facilitating the removal of particles in the luminal space.

## RESULTS

### Renal macrophages make transepithelial protrusions into the tubules

We previously demonstrated that in adult (8~12 weeks old) C57BL/6 mice, the majority of mononuclear phagocytes in the kidneys are F4/80^hi^ MØ which homogenously highly express CX3CR1 ^[27]^. Whole-mount sections of the adult kidneys revealed that F4/80^+^ cells were present in greater density in the medulla relative to the cortex and the papilla (Fig. S1A). Consistently, the kidneys of *Cx3cr1*^GFP/+^ reporter mice showed that GFP^+^ cells were over-represented in the medulla relative to the other anatomical positions, and almost all of them in the medulla were positive for F4/80 (Fig. S1B). In the medulla, we observed a notable spatial proximity of F4/80^+^ cells to renal tubules, as 84.1±0.7% (mean ± SEM) of the F4/80^+^ cells took up residency physically in contact with the tubular epithelium (Fig. 1A). By contrast, the niches encompassing F4/80^+^ cell in the cortex were more diversified: while some F4/80^+^ cells were similarly in the immediate vicinity of tubular epithelium, others were adjacent to the parietal layer of Bowman’s capsule, inside glomerulus, or even in association with sympathetic nerves (Fig. S2A). Many juxtatubular F4/80^+^ cells extended multiple protrusions from their cell bodies in the medulla tissue, whereas the complexity of protrusion in both number and length on per cell basis was less pronounced in the cortical juxtatubular F4/80^+^ cells (Fig. 1B). Such morphological difference could also be observed when GFP^+^ cells in the medulla and in the cortex of *Cx3cr1*^GFP/+^ mice were compared (Fig. S2B).

**Figure 1.**
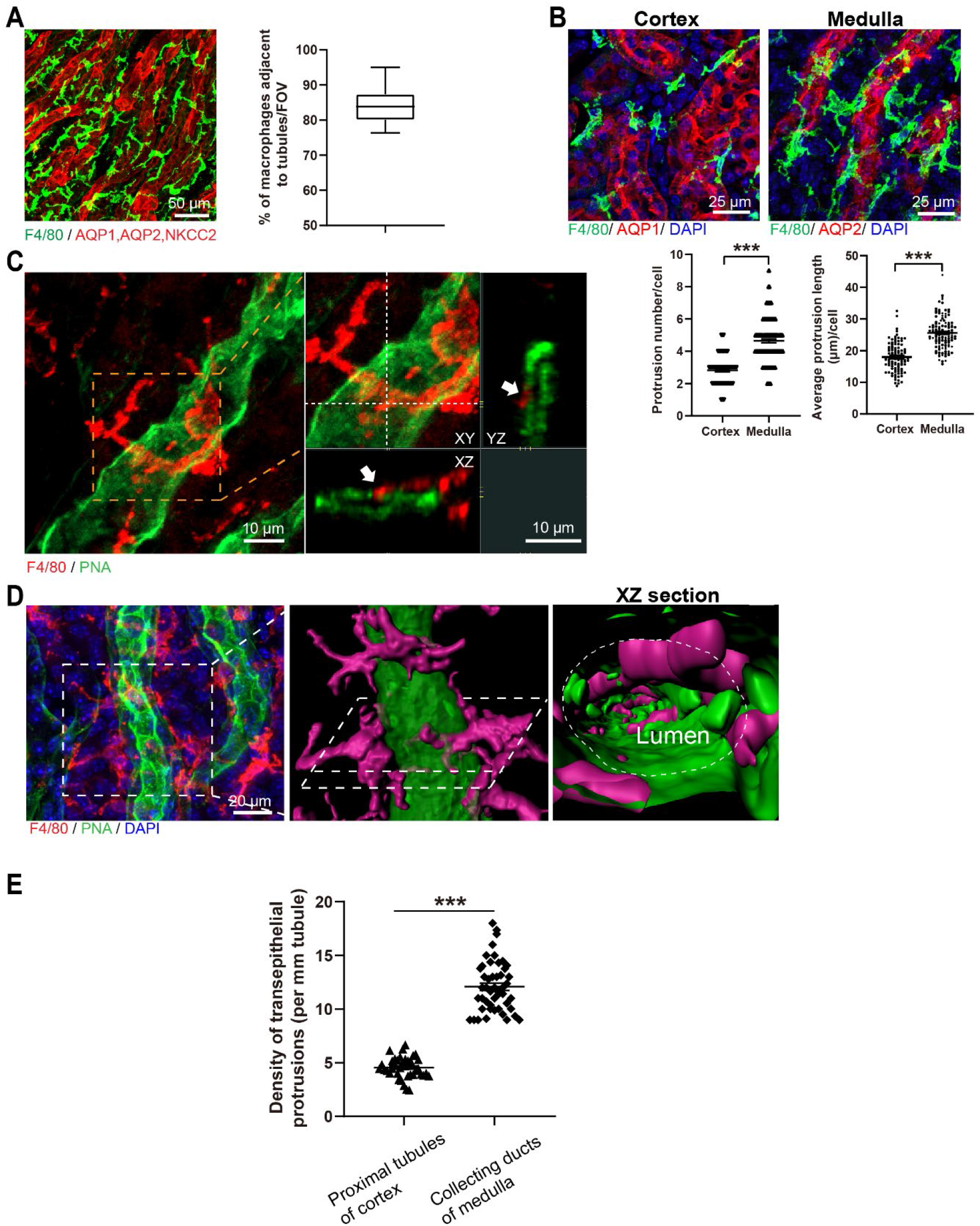
Close interaction between MØ and the medullary collecting ducts. (**A**) Immunofluorescent staining for F4/80 in the renal medulla with z-projection (18 μm). AQP1, AQP2 and NKCC2 together mark all medullary tubules. Quantification of data pooled from 48 fields of view (FOVs) with a total of 3,486 F4/80^+^ cells, shown as a box-and-whiskers graph with 2.5 to 97.5 percentiles. (**B**) (Upper) Confocal images showing morphological differences of F4/80^+^ cells in the juxtatubular zones between the cortex and medulla. Z-projection of 21 μm. AQP1 and AQP2 mark proximal tubules of the cortex and collecting ducts of the medulla, respectively. (Lower) Quantification of protrusion number and average protrusion length per F4/80^+^ cell. n = 6. (**C**) Representative confocal image of medulla sections shows that a part of a protrusion (arrows) derived from a juxtatubular F4/80^+^ cell (red) is exposed to the PNA^+^ apical surface of a collecting duct (green). Z-projections of 24 μm. (**D**) Confocal image of the spatial relationship between PNA^+^ epithelium (green) and F4/80^+^ cell protrusions (red). The magnifications show the corresponding 3D reconstructions viewed from abluminal (left) and luminal (right) sides. (**E**) Quantification of the densities of transepithelial protrusions on the indicated tubule segments. Each dot indicates the quantification from one 319 × 319 μm^2^ FOV. n = 6. \*\*\**P* < 0.005 by two-tailed unpaired t test. Data are depicted as mean ± SEM.

Unexpectedly, confocal analysis of thicker sections (30 μm) of the medulla with high resolution and optical magnification revealed that many of the protrusions derived from F4/80^+^cells penetrated through the laminin^+^ paratubular basement membrane (Fig. S3A) and AQP2^+^ epithelial cells (Fig. S3B), which was confirmed with orthogonal views, corresponding to the ‘‘transepithelial dendrites” of phagocytes previously described in the small intestine ^[28]^. Co-staining F4/80^+^ cells with lectin peanut agglutinin (PNA) which preferentially binds to the apical surface of collecting ducts confirmed that many of the protrusions were transepithelial to the tubular lumen (Fig. 1C). Intraluminal 3D rendering of medullary collecting ducts exhibited protrusion segments inlaying the luminal wall of tubules (Fig. 1D). Systemic analysis revealed that some F4/80^+^ cells could extend more than one transepithelial protrusions (Fig. S3C).

To investigate the possibility that the juxtatubular F4/80^+^ cells we examined had an admixture with dendritic cells (DCs), we employed the reporter *Zbtb46*^GFP/+^ mice in which classical DCs express GFP ^[29]^. We did observe the presence of classical DCs in the renal medulla but they did not overlap with F4/80^+^ cells (Fig. S4A). More importantly, we did not observe transepithelial protrusions derived from DCs (Fig. S4B), suggesting that the transepithelial property in the medulla was exclusive to MØ.

Discrimination with tubular segment-specific markers and anatomical positions showed that the density of MØ transepithelial protrusions in the distal region of nephron, i.e., medullary collecting ducts, was markedly higher than that in the proximal region, i.e., cortical proximal tubules (Fig. 1E). This observation was intriguing because both the transepithelial electrical resistance ^[30]^ and number of tight junction strands ^[31]^ indicated that the medullary collecting tubule had the highest sealing capacity with lowest permeability. To understand how the protrusions of MØ extend through tubular epithelium of collecting ducts, we co-stained F4/80 with tight junctional scaffolding protein ZO-1 (Fig. 2) or adherent junction proteins β-catenin (Fig. S5). This displayed an exclusive expression of junction proteins in the epithelium (not in MØ), resulting in strong and reliable labeling of epithelial cell borders. It revealed that each protrusion (~25 μm in length) in most cases wrapped around more than one epithelial cells (~10 μm in diameter); if the transepithelial part of a protrusion cross over two epithelial cells, it was embedded into epithelial cells almost exclusively through epithelial cell body but circumvented the cell-cell border of epithelium without disrupting the overall junctional architecture (Fig. 2). These observations indicate that MØ protrude through a “transcellular” route instead of between epithelial cells. This is in sharp contrast with intestinal subepithelial phagocytes which inset transepithelial protrusions through lateral “paracellular” space of epithelial cells ^[32, 33]^.

**Figure 2.**
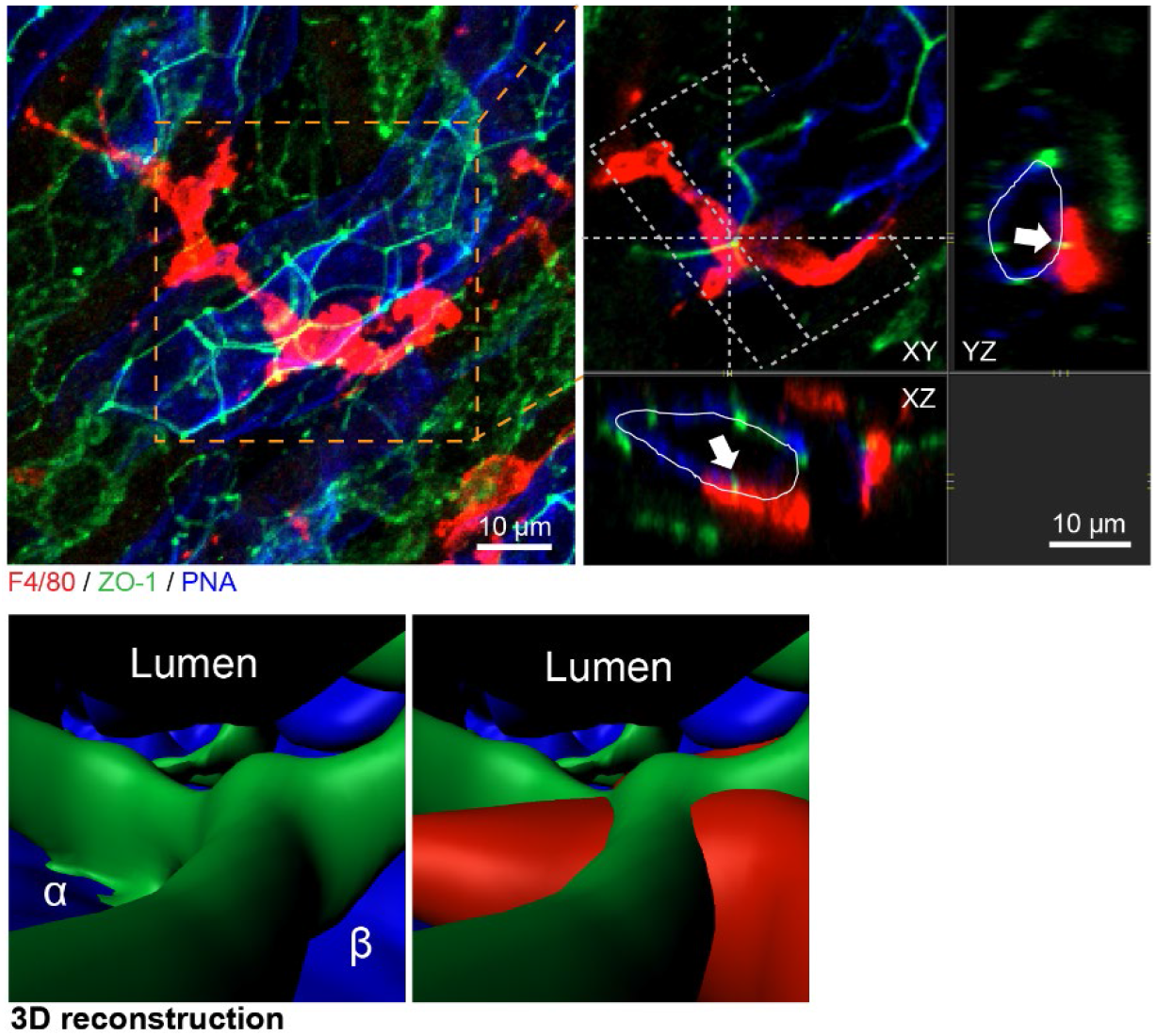
MØ make transepithelial protrusions through a transcellular route. Representative confocal image of a transepithelial protrusion derived from a F4/80^+^ cell (red) with epithelial border protein ZO-1 (green) on the apical surface of a collecting duct (blue). The arrows indicate an intact border structure (green) wrapped around by the protrusion (red). Z-projections of 22 μm. Intraluminal 3D-reconstruction is also shown. α and β, two juxtaposing epithelial cells.

### Medullary juxtatubular macrophages constantly sample urine contents

Next, we were to investigate the behaviors of juxtatubular MØ and potential interactions between MØ and the tubular epithelial barrier. The penetration ability of two-photon excitation microscopy makes it impracticable to detect medulla which lies innermost in the kidney *in vivo*. To circumvent this obstacle, we optimized an approach for live monitoring medulla *ex vivo*. Kidneys from *Cx3cr1*^CreERT2/+^: *R26*^tdTomato^ mice which highlighted MØ after tamoxifen application were gently sliced using a vibrotome in ice-cold buffer to preserve the microarchitecture of the kidney and the viability of cells. To minimally disrupt the structure of kidney, the sliced sections were 200 μm in thickness, much larger than the diameter of collecting ducts which are in a range of 10-20 μm. After warming the sections to 37 °C to restore cell mobility, we tracked the cells by two-photon microscope for up to 2.5 hr in the depth of 100 μm. During the imaging sessions, while the cellular bodies of most juxtatubular MØ remained immobile on epithelial monolayer, their transepithelial protrusions displayed dynamic extension-and-retraction movements, appearing to be sampling the urine contents and probing intratubular environment (Fig. 3A). Accordingly, all (100%) transepithelial protrusions were enriched in LAMP1, a marker of late-endosome/lysosome (Fig. 3B). Using image analysis software to quantify the motion of transepithelial protrusions, we found that, on average, this sampling-like movement per protrusion occurred at a frequency of ~5 times/hr in the medullary collecting ducts, over twofold faster than the similar movement of MØ protrusions inside the proximal epithelium of cortex (Fig. 3C).

**Figure 3.**
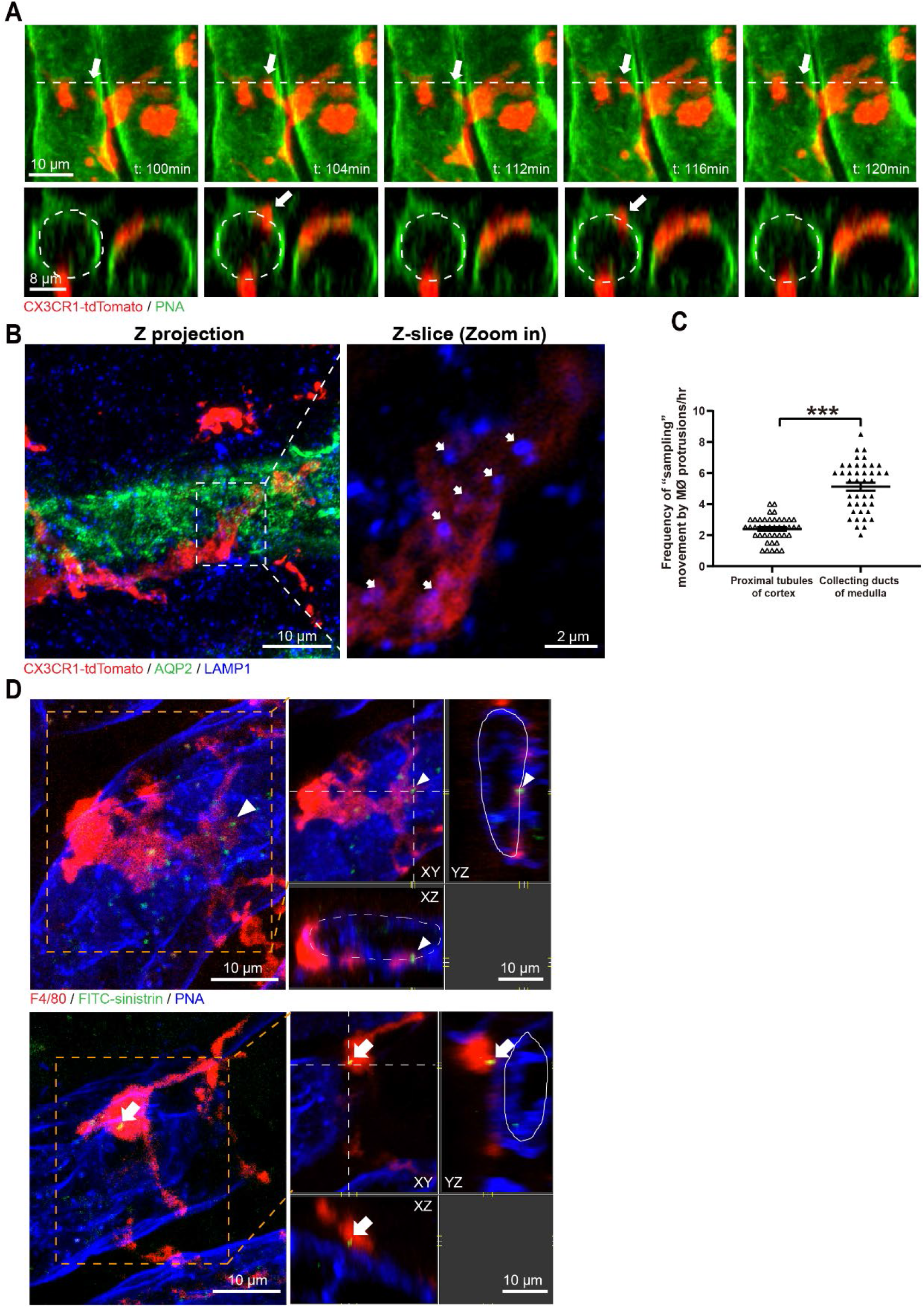
Medullary juxtatubular macrophages are sampling urine contents through their transepithelial protrusions. (**A**) Time lapse of a transepithelial protrusion (arrow) of tdTomato^+^ MØ (red) that makes repetitive extension-and-retraction movements inside a medullary collecting duct (green). The border of tubular lumen is indicated with dashed line. Z-projections of 30 μm. (**B**) The LAMP1-enriched late-endosomes/lysosomes (blue) distribute in the MØ transepithelial protrusions (red). AQP2 indicates the epithelial cells of medullary collecting ducts. Z-projections of 27 μm. (**C**) Quantification of the frequencies of the sampling-like movement of MØ transepithelial protrusions inside cortical proximal tubules (n = 6) and medullary collecting ducts (n = 7). \*\*\**P* < 0.005 by two-tailed unpaired t test. Data are depicted as mean ± SEM. (**D**) The juxtatubular MØ and their transepithelial protrusions (red) in the medullary collecting ducts (green) were examined 60 min post i.v. infusion of FITC-sinistrin. Z-projections of 21 μm. Note that by 60 min, a transepithelial protrusion (arrowhead of the left panel) of MØ and a MØ soma (arrow of the right panel) outside of the tubules had contained FITC-sinistrin particles.

To probe the hypothesis that the sampling-like movement of juxtatubular MØ indeed reflects dynamic intake of intratubular materials, we *i.v*. injected fluorescein isothiocyanate (FITC)-labeled sinistrin which was widely used for measuring glomerular filtration rate (GFR) since it will be quickly and exclusively excreted through urine ^[34]^. Sinistrin rapidly emerged in the collecting ducts within 30 min post *i.v*. infusion (Fig. S6A), and disappeared rapidly from the skin vasculature (within 60 min) (Fig. S6B), consistent with glomerular filtration. By 60 min, 62.9 ± 1.9% (mean ± SEM) of the MØ transepithelial protrusions in the medullary collecting duct, as well as the cell bodies of some MØ, had contained FITC-sinistrin (Fig. 3D). In this time frame, intake of FITC-sinistrin by epithelial cells was not noticed, supporting a role of the transepithelial protrusions in sampling urine contents.

### A critical role of medulla macrophages in disposing urine particles

The observed behavior of MØ in sampling urine contents provoked us to interrogate the potential interactions of juxtatubular MØ with big particles in the urine, since natural particles are prone to form and accumulate at the caudal segment of nephron, i.e., medullary collecting ducts. To this end, we performed intrapelvic injection of fluorescent inert latex beads (2 μm in diameter) which do not attract immune cells via chemotaxis. Two hours after administration, beads could be conspicuously observed in the medullary collecting ducts but not in the proximal tubules (Fig. S7A). At this time point, about 52.4 ± 2.3% (mean ± SEM) of beads had already been associated with MØ transepithelial protrusions (Fig. 4A). By 12 hours, some beads had been intracellularly transported to the MØ soma positioned at the basolateral side of tubules (Fig. 4B); we could occasionally observe the accumulation of multiple beads in one MØ cell body (Fig. S7B). Indeed, by 12 hours, free beads were almost cleared off from the lumen of the collecting ducts (Fig. 4C), suggesting an extremely effective particle sequestration executed by MØ in addition to urine flush. To corroborate the role of MØ in particle disposal, we depleted renal MØ with liposome-clodronate before intrapelvic loading of beads. When the kidneys were examined 36 hr post bead injection, a dramatic increase of bead retention in the kidney was noticed in the MØ-depleted mice compared to the liposome-PBS treated MØ-intact controls (Fig. 4D), suggesting that urine flushing was not sufficient to remove big particles in the renal tubule system without the assistance of MØ-mediated disposal.

**Figure 4.**
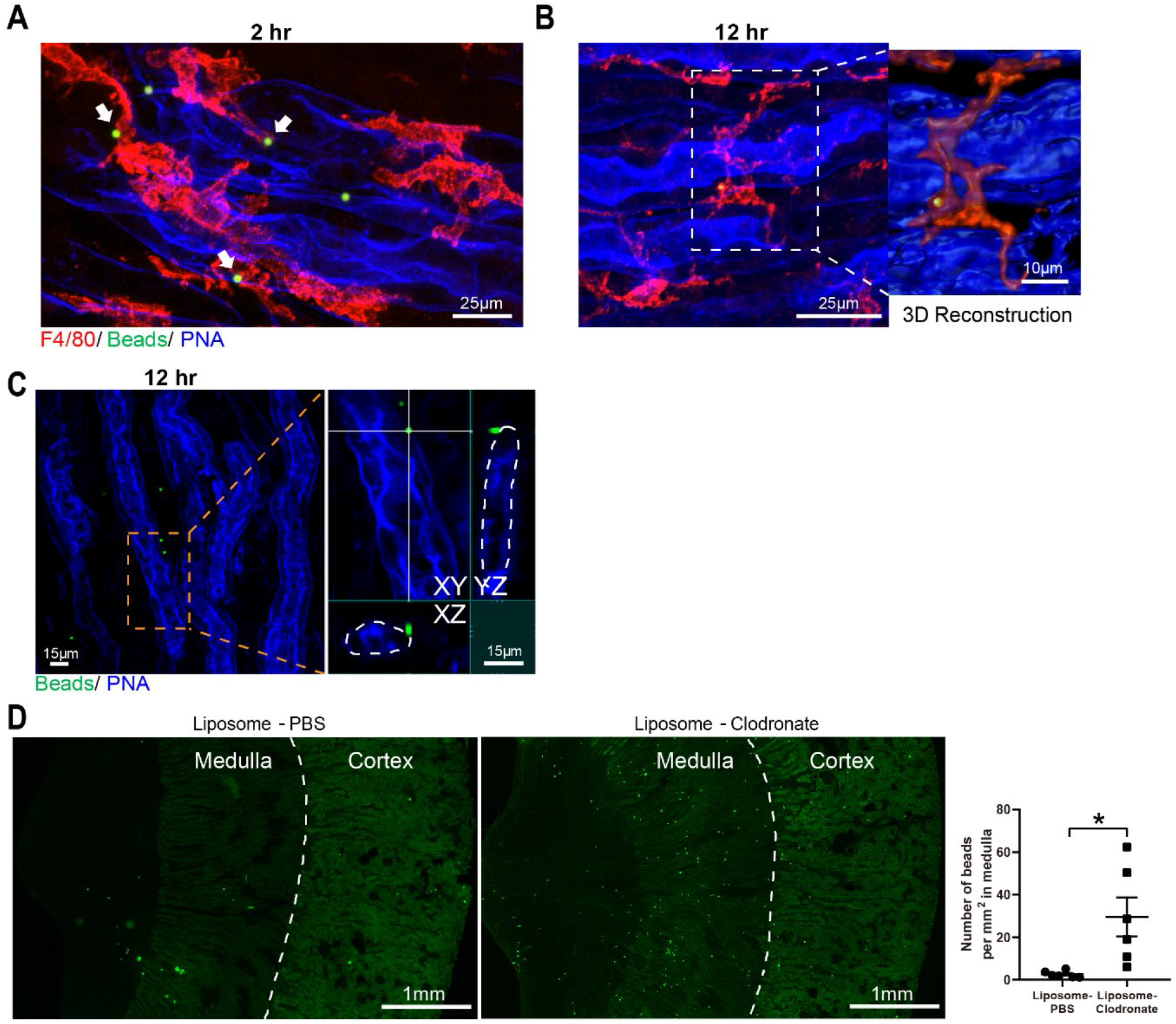
A role of medullary juxtatubular macrophages in disposing urine particles. (**A**, **B**) Mice received intrapelvic injection of fluorescent latex beads (2 μm in diameter). After 2 hr (A) and 12 hr (B), the interactions between MØ (red) and beads (green) were examined around medullary collecting ducts (blue). Arrow, MØ protrusion-associated bead. z-projection of 23 μm (A). A bead in the soma of a juxtatubular MØ; z-projection of 20 μm; 3D-reconstruction is shown in the magnified inset (B). (**C**) Twelve hours post bead administration, beads were almost absent in the intratubular compartment. The dashed line in the magnified inset indicates the border of tubular lumen. (**D**) Mice were first i.p. treated with either liposome-PBS or liposome-clodronate twice with 24 hr apart. Twenty-four hours post the last treatment, fluorescent latex beads were intrapelvically injected and the kidneys were examined 36 hr later. Representative pictures and the quantification of bead densities in the medulla are shown. n = 6. \**P* < 0.05 by two-tailed unpaired t test. Data are depicted as mean ± SEM.

Albeit moving particles to the abluminal side is an efficient tactic to unblock the tubules, we suspected that there was other way by which MØ dispose intratubular particles in addition to the lumen-to-interstitium transportation, since we did not observe an accumulation of latex beads in the renal interstitium in the MØ-intact mice (Liposome-PBS group, Fig. 4D). We hypothesized that MØ could gain access to the lumen and “escort” particles to be discharged via urine. To probe this possibility, we first assessed whether there were transmigrated MØ in the urine of naïve mice. To preclude contaminants from ureter and bladder, we directly collected urine via a transpelvic route. Consistent with our hypothesis, F4/80^hi^ MØ were indeed present in the normal urine and they accounted for the majority of CD45^+^ immune cells in the urine (Fig. 5A). F4/80^hi^ cells were absent in the blood, excluding blood contamination (Fig. S8). Almost all of the urinary F4/80^hi^ cells appeared high in forward scatter area (FSC-A) when examined by flow cytometry (Fig. 5A), suggesting their association with urine particles. Indeed, a significant portion of F4/80^hi^ cells were also positive for AQP2 (Fig. 5A), a marker of collecting duct epithelial cells, indicative of an association between MØ and sloughed epithelial cells. Supporting the hypothesis that “escorting” is another strategy employed by MØ to remove urinary particles, we could detect bead-bearing MØ in the urine post intrapelvic latex bead injection (Fig. 5B).

**Figure 5.**
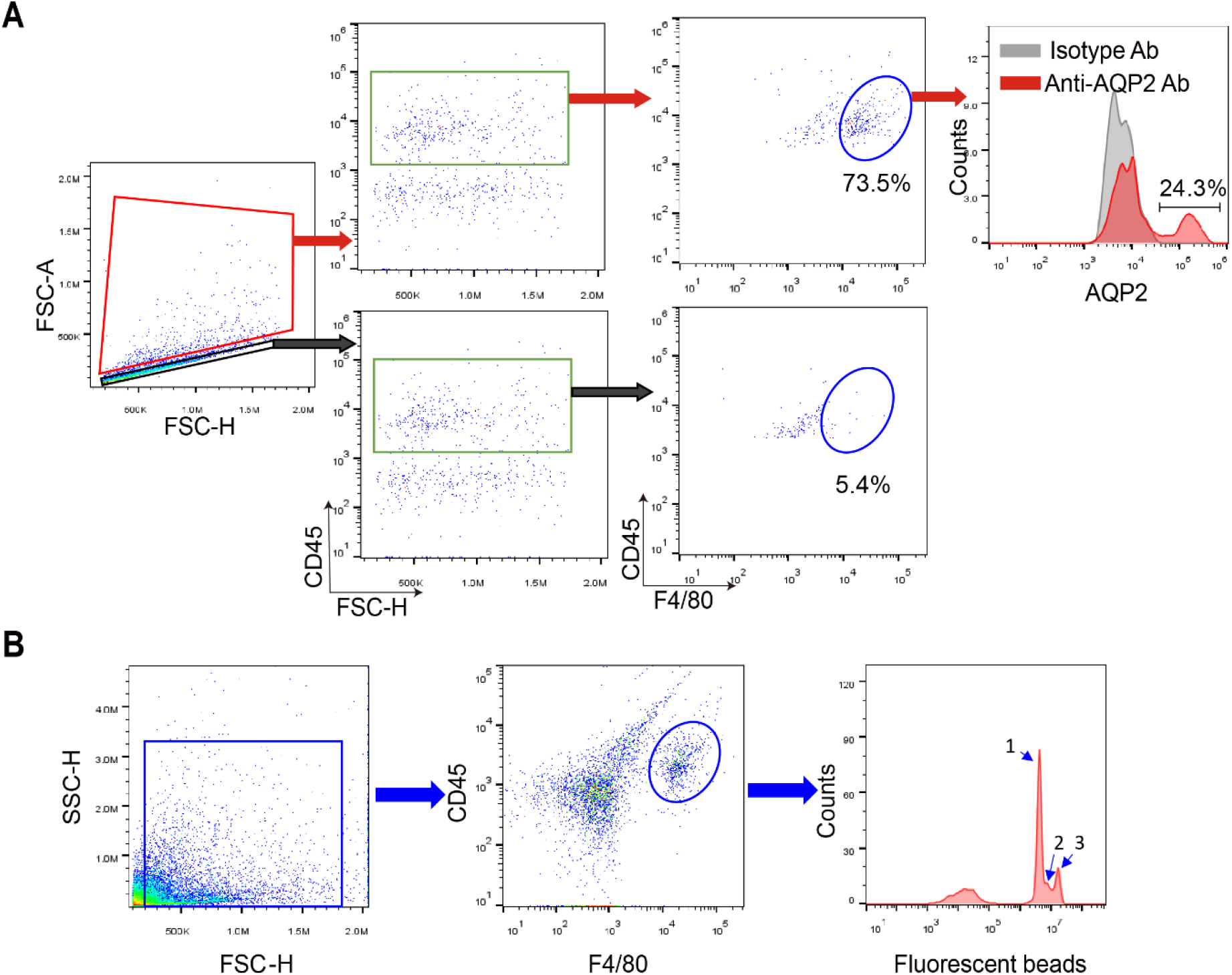
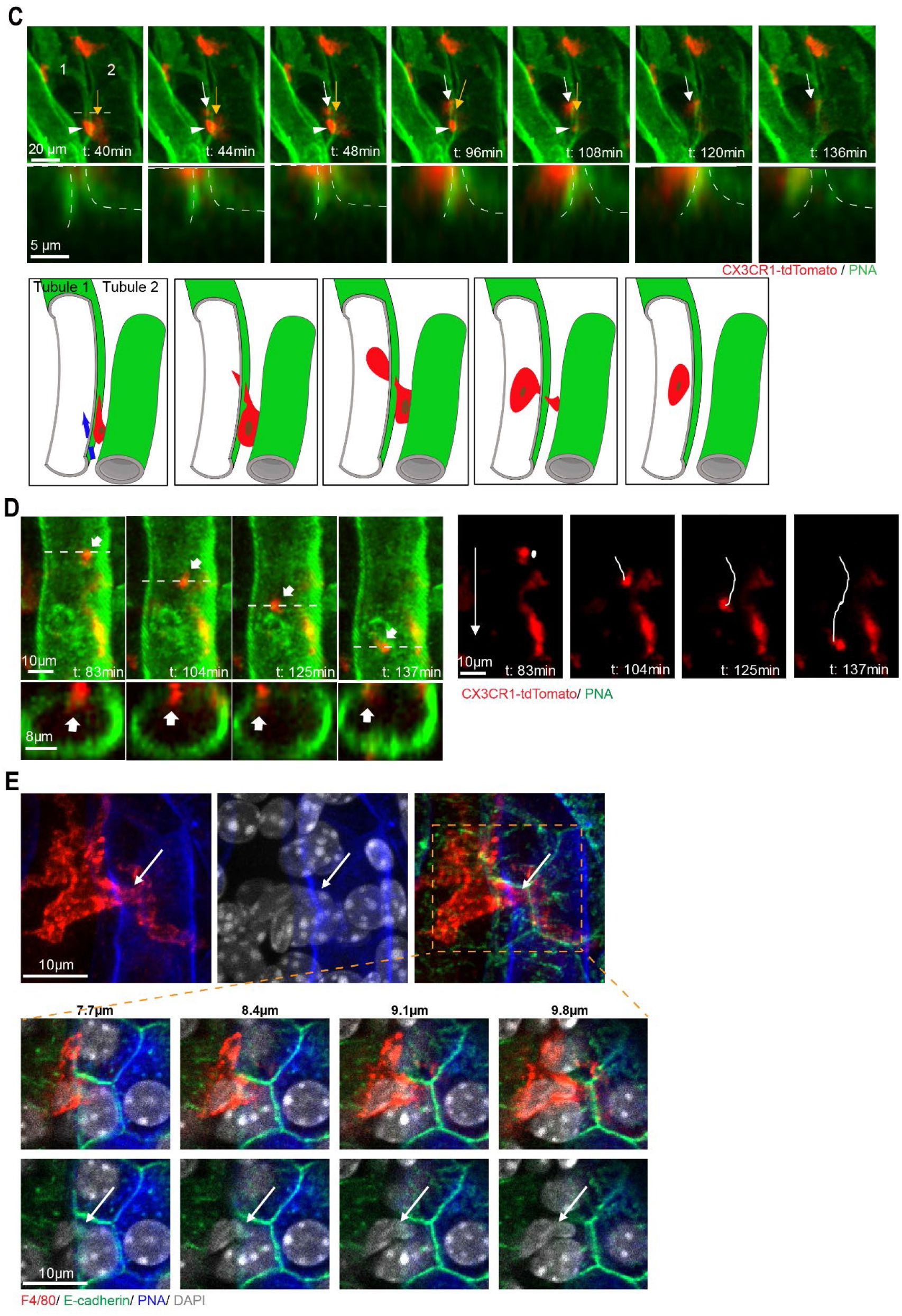
Medullary juxtatubular macrophages can make transepithelial migration. Representative flow cytometry analysis of the urine collected from the pelvis of naive C57BL/6 mice. Most F4/80^hi^ MØ are in the FSC-A^hi^ zone and they account for the majority of CD45^+^ immune cells in the urine. AQP2, a signature molecule of collecting duct epithelial cells, could be co-stained in a portion of MØ. (**B**) Four hours after intrapelvic administration of fluorescent latex beads, urine was collected and subject to flow cytometry analysis which shows that the majority of CD45^+^F4/80^hi^ MØ are associated with beads. The numbers on the graph, i.e., “1”, “2” and “3”, indicate the cells in association with the corresponding number of beads on per cell basis. (**C**) Time lapse of a MØ (red) making transepithelial migration to the lumen of a collecting duct (green) in the medulla. Arrowhead, the cell body in the interstitium between two collecting ducts; yellow arrow, the squeezed part of the cell when transmigrating into the epithelium of the tubule left to the cell, white arrow, the transmigrated cell part present in the tubular lumen. The arrow in the cartoon shows direction of movement. (**D**) Time-lapse image sequence of a MØ (red, indicated by an arrow in the left panel) rolling inside the tubular lumen of a collecting duct (green) in the medulla. The arrow in the right panels denotes the path of the MØ, with the track in white. (**E**) Immunofluorescent staining shows a MØ (red) with its nucleus (white) inserted into the epithelial monolayer (blue, with epithelial border denoted in green) in the renal medulla. Upper panels, z-projection of 17 μm; Lower panels, individual optical planes depicting the same MØ at different depths.

Consistent with the appearance of MØ in the urine, two-photon microscopy of live kidney sections unveiled that juxtatubular MØ occasionally made transepithelial migration into the lumen of medullary collecting ducts (Fig. 5C). This phenomenon was only observed in the slices of medulla but not those of cortex. The still images (orthogonal cross-sections through the 3D stacks in successive time frames) of a real-time video showed that at the beginning, the cell body of a fluorescent MØ sat in the interstitium between two collecting ducts with a preformed protrusion inserting into the epithelium of one adjacent tubule. From then on, the cell body started to squeeze into the transepithelial channel made by the protrusion till completion of migration (Fig. 5C). Afterwards, the migrated MØ moved away from the entry site in the tubular lumen. The transepithelial migration was consistent among more than 12 mice and 24 videos analyzed. Moreover, we observed crawling MØ which lost polarity and showed a sphere-like shape inside collecting ducts (Fig. 5D). Further supporting the capability of juxtatubular MØ in making transepithelial migration, confocal microscopy of fixed slides could identify that some MØ inserted part of their nucleus into the epithelial monolayer (Fig. 5E), which was often observed when transendothelial migration of leukocytes take place under inflammation ^[35]^. These data, together with the observation of transepithelial protrusions formed by renal MØ, resonate with a recently emerging finding that protrusion insertion is a complementary mechanism to allow cell migration without specific adhesions ^[36]^.

### Renal macrophages protect against kidney stone formation and related inflammation

The aforementioned role of MØ in sampling and disposing urine particles was studied in the context of a nonpathogenic reagent. Physiologically, kidneys are continuously exposed to mineral crystallization because of urine supersaturation, a situation particularly concerning medullary collecting tubule which is the caudal segment of the renal tubular system. Calcium oxalate stone is the most common type of nephrolithiasis ^[37]^. To gain insights into the physiological impact of juxtatubular MØ on the development of nephrolithiasis, we induced an acute hyperoxaluria model by treating C57BL/6 mice with a single intraperitoneal (i.p.) injection of sodium oxalate (NaOx) (100 mg/kg) plus 3% NaOx in drinking water. This challenge led to rapid, abundant CaOx crystal deposition exclusively luminal in 24 hr (Fig. S9A). Crystals were mostly located in the medulla, which is in line with the fact that the urine concentration mainly occurs in the medulla. The crystals frequently caused obstruction, as revealed by tubule dilation (Fig. S9B). Many crystals at this moment were surrounded by MØ in the lumen, appearing being phagocytosed (Fig. 6A). Interestingly, CaOx crystallization was accompanied with a marked increase of the number of transepithelial protrusions extended by MØ in the medullary collecting ducts (Fig. 6B). Moreover, live recording revealed that the protrusions of MØ in the collecting ducts enhanced motility, indicating an expedited “sampling” behavior (Fig. 6C).

**Figure 6.**
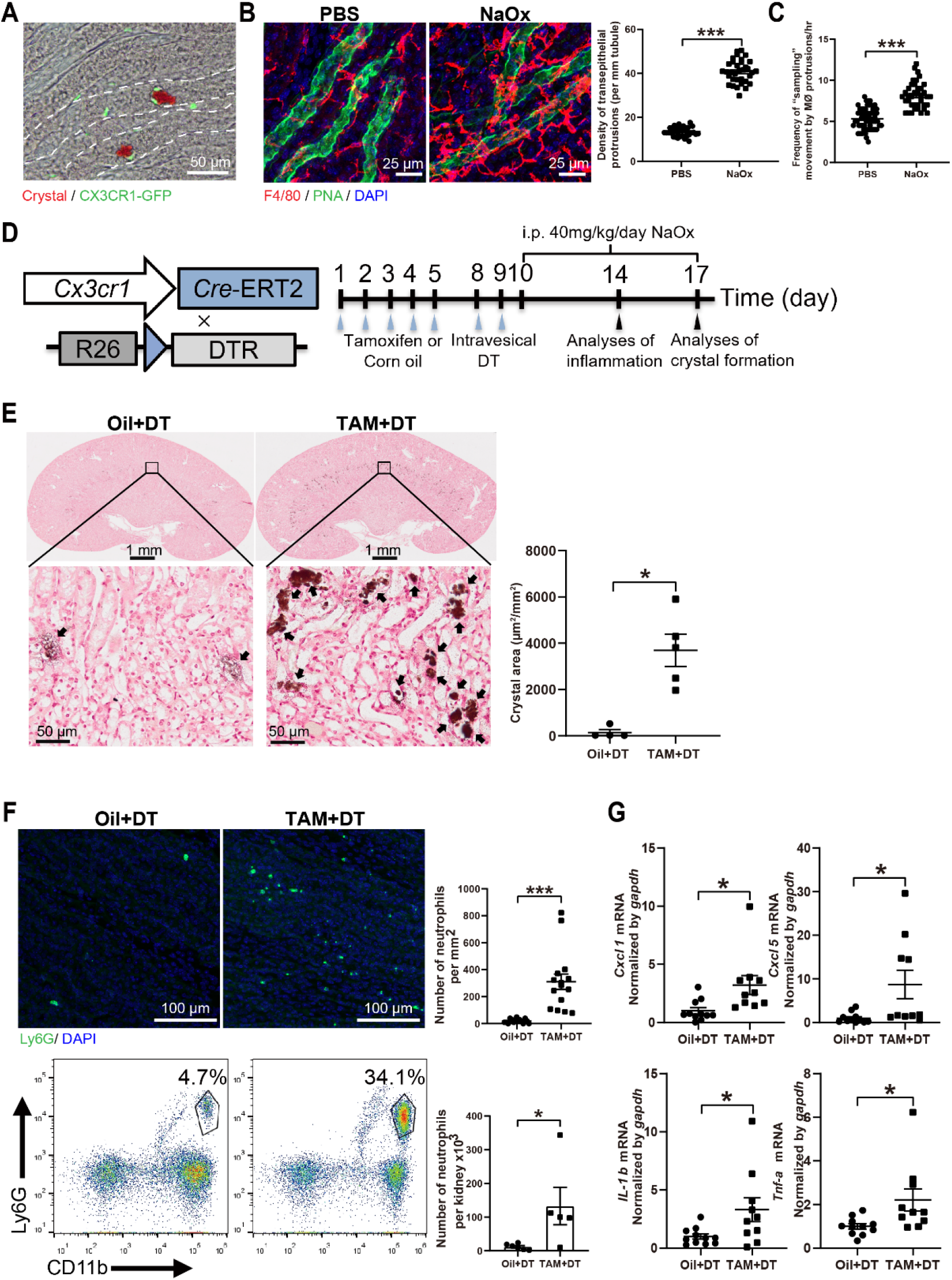
Renal macrophages protect against kidney stone formation and related inflammation. (**A-C**) In an acute hyperoxaluria model, mice were challenged with i.p. injection of NaOx (100 mg/kg) plus drinking 3% NaOx water. The mice were examined 24 hr later. The controls were i.p. treated with PBS and drank normal water. (**A**) Alizarin red S staining of the medulla of *Cx3cr1*^GFP/+^ mice shows that MØ (green) were intimately associated with CaOx crystals (red) in the tubules. The dashed lines delineate the contour of the tubules. (**B**) Confocal images of MØ (red) and the collecting ducts (green) in the medulla after the indicated treatments, with z-projection of 19 μm. Quantification of the densities of the transepithelial protrusions was also shown. n = 6. (**C**) Quantification of the frequency of the sampling-like movement by MØ transepithelial protrusions in the medullary collecting ducts after the indicated treatments. n = 4. (**D**) Scheme illustrating kidney-specific MØ depletion. *Cx3cr1^CreER/+^:Rosa26-iDTR* mice were treated with either corn oil (as controls) or tamoxifen (TAM, for irreversible expression of DTR in CX3CR1^+^ cells), followed by two injections of DT 24 hr apart. Thereafter, a chronic hyperoxaluria model was followed. (**E**-**G**) Experiments were performed with the protocol shown in **D**. (**E**) Seven days after the onset of NaOx treatment, CaOx crystal deposition was examined by Von Kossa staining. Quantification of the affected area in the medulla was also shown. Oil+DT, n = 4; TAM+DT, n = 5. (**F, G**) Four days after the onset of NaOx treatment, the number of intrarenal neutrophils was quantitated by both immunohistostaining of Ly6G (green) (the upper panels, n = 3) and flow cytometry analysis of CD45^+^ cells (the lower panels; Oil+DT, n = 6; TAM+DT, n = 5). (**F**); relative expression of the indicated inflammatory cytokines/chemokines in the kidneys, assessed by quantitative RT-PCR (Oil+DT, n = 11; TAM+DT, n = 10). (**G**). \**P* < 0.05, \*\*\**P* < 0.005 by two-tailed unpaired t test. Data are depicted as mean ± SEM.

To examine a role of renal MØ, we challenged mice with a chronic hyperoxaluria model in which CaOx crystallization develops in a slower pattern resembling a process taking place in nephrolithiasis patients. By doing that, we i.p. treated mice daily with a low dose of NaOx (40 mg/kg) for 7 days without additional NaOx drinking, resulting in a mild renal CaOx crystallization in adult C57BL/6 mice (Fig. S9C). To selectively remove renal MØ, we crossed *Cx3cr1*^CreERT2^ mice to *Rosa26-iDTR* mice. After their female progeny *Cx3cr1^CreER/+^:iDTR* mice were treated with tamoxifen for inducing diphtheria toxin receptor (DTR) expression on renal MØ, we adopted an intravesical DT application strategy (see Methods) (Fig. 6D). This strategy was successful in removing renal medullary MØ (Fig. S10A), sparing blood monocytes (Fig. S10B) and brain microglia (Fig. S10C) which also express CX3CR1 but reside in anatomically disparate compartments. Loss of medullary MØ (TAM+DT) during hyperoxaluria challenge significantly precipitated kidney stone formation in comparison to the MØ-intact controls (Oil+DT) under the same insult, as the affected renal area expanded by 25.4 folds (Fig. 6E), suggesting a critical role of renal MØ in preventing kidney stone formation. Nephrolithiasis is accompanied with local inflammation ^[37]^. We noticed that there were significantly more neutrophils infiltrating into the kidneys of MØ-depleted mice relative to MØ-intact mice during hyperoxaluria challenge, as assessed by both immunofluorescent staining and flow cytometry (Fig. 6F). In line, MØ-depleted mice exhibited markedly higher levels of intrarenal chemokines and pro-inflammatory cytokines than their MØ-intact littermates during hyperoxaluria challenge (Fig. 6G). To be noted, MØ depletion itself did not cause an overt inflammation (Figs. S11A and S11B). These data suggest that aggravated crystal-induced tissue disruption in the absence of resident medullary MØ resulted in greater inflammation. These observations highlight the role of MØ as an effective guardian that precludes both crystal deposition and possible damages inflicted by secondary inflammation.

## DISCUSSION

During the filtrate of the glomerulus flows though the renal tubular system, sloughed tubular epithelial cells and mineral crystals resulting from urine concentration generate numerous microscopic sediment particles. In pathological conditions, the urine sediment would also include aggregates of pathogens, large-size proteins such as antibodies, and even inflammatory cells ^[38–40]^. Lodging of these particles in the tubules will eventually block the passage of urine and lead to renal dysfunction. Thus, it is critical to forestall an obstruction of the affected tubules for maintaining the free flow of urine; however, the underlying mechanisms stay unexplored. The present study uncovers that renal MØ, in particular those juxtaposing to the medullary collecting ducts, are dedicated to removing particles in the tubule system. These phagocytes extend transepithelial protrusion and constitutively uptake urine materials. In particular, they are proficient in sequestering and phagocytosing intraluminal particles. Moreover, the juxtatubular MØ occasionally make transmigration to the tubule lumen to escort the excretion of urine particles. This unique behavior is live recorded by two-photon microscopy, and is also manifested by the detection of insertion of MØ nucleus into the epithelial monolayer and detection of MØ presence in the normal urine. The importance of the roles of renal MØ in dislodging tubular sediments was evidenced by both a significant delay of the removal of intratubular beads and a precipitation of nephrolithiasis upon oxalate salt challenge when devoid of renal MØ. The physiological significance of this MØ-associated function was further highlighted by the longitudinal studies of MØ underrepresentation models: by both genetic and pharmacological approaches, we demonstrated that a long-term renal MØ deficiency would result in rampant accumulation of diverse urine sediments in the tubules. Thus, our data imply that urine flushing alone is not enough to keep the renal tubules unobstructed, but a routine “plumber” work executed by MØ is additionally required.

Our time-lapse two-photon microscope observation display that the transepithelial protrusions of juxtatubular MØ routinely make continuous extension-retraction movements. More importantly, through these sampling-like movements, MØ take up urine materials, including large-size particles. The juxtatubular MØ are ideally located to provide surveillance of epithelial cells and intratubular environments. In response to urinary irritants e.g., mineral crystals, these MØ could enhance the motility of their protrusions to expedite removal of the intratubular irritants. Thus these phagocytes in this particular position serve as a “gatekeeper” function within the complex structure of the kidney, predicting that the juxtatubular MØ are the first immune cell type to interact with substances and infectious agents entering the kidney through the urinary tract. Conceivably, the particular position and special behavior of juxtatubular MØ could digest and present antigens derived from pathogens retrogradely disseminated from downstream urinary tract. With this perspective, further exploration of their defensive roles is warranted.

Of note, MØ were found to be the major immune cells present in the normal urine collected from the pelvis, indicating that MØ have the ability to gain access into the intraluminal compartment of the kidney. Supporting an escort role of MØ in removing tubular deposits, the urinary MØ were generally associated with urine particles including epithelial cell fragments and the injected microspheres. Leukocyte extravasation has been studied for decades and the pertinent machinery of the interaction between endothelial and leukocytes and the followed leukocyte vascular egress has been well described ^[41, 42]^. However, crossing the epithelial barrier is in a basolateral-to-apical direction, opposite to diapedesis. The few membrane proteins proposed to regulate transepithelial migration have largely been extrapolated from the studies of intestinal and pulmonary epithelium ^[33, 43–45]^, and many such studies were performed in the context of inflammation. Due to the complexity of cellular pseudopodia formation and of the interaction between phagocytes and epithelial cells in varying context, defining the dynamics of MØ-epithelial interactions and the related molecules involved during the formation of transepithelial protrusions in the future is necessary for better understating renal architecture the interplay between resident tissue cells.

In an anatomical perspective, we noticed an intriguing disparity between cortical and medullary juxtatubular MØ in their morphology and transepithelial behaviors, with more pronounced transepithelial protrusions with the medullary MØ. It is well appreciated that the microenvironments between cortex and medulla are very different in that the medulla is hyperosmotic and hypoxic ^[4, 46–48]^. Especially, hypersalinity was shown to induce epithelial cells to secrete chemokines for recruiting monocyte-derived phagocytes to the renal medulla, where this hypersaline environment also increased the intrinsic bactericidal of these phagocytes ^[49]^. Whether these milieu cues also drive the morphological and functional characteristics of juxtatubular MØ deserves further exploration.

Taken together, our findings suggest that renal MØ, heretofore primarily considered in the context of immune responses and tissue remodeling, play an important role in monitoring urine contents and facilitating removal of urine sedimentary particles. A potential utilization of this distinctive feature of renal MØ may be beneficial for developing therapeutics for nephrolithiasis and kidney-specific drug delivery.

## Materials and Methods

### Mice

*Cx3cr1*^GFP/+^ mice [005582], *Zbtb46*^GFP/+^ mice [027618]*, Cx3cr1*^CreERT2^ mice [020940], *Rosa26*-iDTR mice [007900] and *Rosa26-stop-TdTomato* mice [007914] were from The Jackson Laboratory. All the mice are in C57BL/6 background. Normal C57BL/6 mice (CD45.2^+^) were purchased from Shanghai Research Center for Model Organisms. All mice used in this study without specific explanation were 8- to 12-week-old males. Mice were housed in a standard animal facility, with a 12-hr light/dark cycle, in specific-pathogen-free environment. All animal experiments were approved by the Institutional Animal Care and Use Committee at Zhejiang University.

### Antibodies

The following fluorophore-conjugated antibodies used for flow cytometry analysis were purchased from BioLegend or Thermo Fisher Scientific: anti-CD3 (145-2C11), anti-CD11b (M1/70), anti-CD19 (6D5), anti-CD45 (30-F11), anti-CD49b (DX5), anti-CD64 (X54-5/7.1), anti-CD115 (AFS98), anti-F4/80 (BM8), anti-CX3CR1 (SA011F11), and anti-Ly6G (1A8). Anti-AQP2 (E-2) antibodies were purchased from R&D and Santa Cruz Biotechnology, respectively. The following antibodies were used for immunohistostaining: anti-laminin and anti-AQP1 were purchased from Sigma-Aldrich; anti-F4/80 and anti-β-catenin were purchased from Thermo Fisher Scientific; anti-ZO-1 was from Proteintech; anti-NKCC2, anti-LAMP1 and anti-GFP were purchased from Abcam; Peanut Agglutinin (PNA) and Lotus Tetragonolobus Lectin (LTL) were purchased from Vector Laboratories; anti-Tdtomato was purchased from Biorbyt.

### Reagents and their *in vivo* treatments

To monitor the interaction between renal MØ with urine contents, FITC-sinistrin (MediBeacon, Mannheim, Germany) was injected *i.v*. at a dose of 7mg/100g. To monitor the interaction between renal MØ with intratubular large particles, the stock 2 μm fluorescent latex beads (2.5%, Sigma) was first diluted 10 times by PBS, and then 25 μl bead solution were injected to the intrapelvic space of anesthetized mice.

### Intrapelvic injection and collection of urine contents

Mice were anesthetized and kept on a homeothermic pad to maintain body temperature during the procedure. Dorsal incision was made to expose kidney. The adipose tissue surrounding the kidney was gently separated to expose the pelvis which locates in the renal hilus. For intrapelvic injection, 25 μl 0.25% fluorescent latex beads (2 μm in diameter) in PBS was injected to the pelvis by aa insulin syringe with a 30 gauge needle. The needle was kept in place for 10 seconds to forestall backflow. After that, the needle was slowly pulled back. The kidney was then returned to the abdominal cavity. For sampling urine contents from the pelvis, 25 μl PBS was first infused into the pelvis followed by gently withdrawing all collectable intrapelvic liquid (~10 μl) using the same insulin syringe.

### Immunostaining and confocal imaging

Mice were anesthetized and transcardially perfused consecutively with cold PBS for 5 min and 4% paraformaldehyde (PFA) for another 5 min. Kidneys were then removed, fixed overnight in 4% PFA, and then dehydrated in PBS containing 30% sucrose at 4°C for 24 hr. Frozen sections (30 μm thick) were made on a cryostat (CM3050S; Leica) at −20 °C. Sections were blocked for 2 hr at room temperature in a blocking buffer (5% donkey serum in PBS). The sections were then incubated with primary antibodies in antibody dilution buffer (0.3% Triton X-100 and 1% BSA in PBS) overnight at 4 °C, followed by incubation with secondary donkey antibodies in antibody dilution buffer for 2 hr at room temperature in the dark. After mounting on glass slides, stained sections were viewed under confocal microscope equipped with 10x /0.45, 20x /0.8, 40x /0.95 NA Plan-Apochromat objective (Carl Zeiss LSM 900). Images were analyzed with the IMARIS software (Bitplane, Switzerland).

### Super-resolution imaging

For imaging the transepithelial protrusions of MØ, super-resolution images of F4/80-labeled MØ and AQP2-labeled collecting duct epithelium were taken by Zeiss LSM 900 Axio Observer confocal microscope with Airyscan 2 system under a Plan-Apochromat 63×1.4 NA objective lens. The Z-resolution was 0.4 um. For these acquisitions, we performed oversampling (pixel size of 10 nm (x, y dimensions)) on the zoomed fragment of the tissue.

### Two-photon imaging of live kidney sections

Two-photon imaging of live kidney sections was developed as a technique for visualizing tissue architecture and cell mobility at close to physiological conditions. Mice were euthanized and the kidneys were harvested and kept on ice in a typical pre-oxygenated bath solution (150 mM NaCl, 5 mM KCl, 1 mM CaCl_2_, 2 mM MgCl_2_, 5 mM glucose, and 10 mM HEPES). The kidneys derived from *Cx3cr1*^CreER/+^: *R26*^tdTomato^ mice were sliced into 200 μm sections with a Leica VT1000 S vibrating blade microtome (Leica Microsystems) at speed 1.80 and frequency 7. Tissue sections were then stained with 0.25 mg/ml PNA in bath solution at 4°C for 30 min, followed by culturing in complete kidney medium (pre-oxygenated phenol-red-free DMEM containing 10% heat-inactivated FBS and 10 mM HEPES) in humidified incubator at 37° C for 2 hr. Sections were held down with tissue anchors in 35 mm glass dishes and were imaged by Olympus FVMPE-RS two-photon microscope equipped with an environmental chamber and a motorized stage. Microscope configuration was set up for four-dimensional analysis (x, y, z, t). The fields were simultaneously recorded with an XLPlan N 25×/1.05 water immersion objective lens every 4 min for 2.5 hr with 1.5 μm Z axis increments and 800×800 pixel resolution. The range of Z stack of images was 25-30 μm. The laser was tuned to 920-nm excitation and used for all studies. Images and videos were processed using Imaris (Bitplane) software.

### Renal immune cell extraction and flow cytometry

Kidneys were isolated from mice after perfusion with cold PBS plus EDTA and were cut into small pieces. Each kidney was digested in 6 ml RPMI1640 (GIBICO) with 1.5 mg/ml collagenase IV (Worthington) and 5 μl/ml DNase I (Sigma) for 30 min at 37°C with gentle shaking. After digestion, cells were passed through a 70-μm strainer (BD) and were then subject to gradient centrifugation by using 72% and 36% Percoll (GE Healthcare). Immune cells were enriched at the 72%/36% interface and were collected for flow cytometry analyses. Samples were analyzed with a 3-laser flow cytometer (Agilent Novocyte) and data were processed with FlowJo (v10.1). Renal macrophages were purified by cell sorting with a BD SORP ARIA II.

### Renal macrophage depletion

For selectively depleting renal MØ, female *Cx3cr1^CreER/+^:iDTR* mice were first treated *i.p*. with tamoxifen (MCE) dissolved in corn oil at a dose of 75 μg/g for 5 consecutive days. After that, mice were intravesically given 2 doses of DT on Day 7 and Day 8. To do so, mice were anaesthetized and intravesically inserted with a PE-10 tubing catheter (RWD Life Science 62324, 0.28 mm × OD 0.61 mm) through which 40 ng/g of DT (Bio Academia) in 80 μl saline was infused. The mice were then placed in a position with their posterior limbs up at a 30-degree angle for 2 hr after the treatment. To evaluate the bead retention in the absence of MØ, mice were injected *i.v*. with 10 μl/g liposome-clodronate (Liposoma) once a day for two days prior to experiment.

### Sodium oxalate-induced calcium oxalate kidney stone model

For developing a rapid kidney stone formation, mice received a single *i.p*. injection of 100 mg/kg NaOx (Sigma) plus 3% NaOx in drinking water. After 24 hr, mice were harvested and the kidneys were examined. Alternatively, we induced a slow crystal deposition model by treating mice *i.p*. with NaOx in a daily dose of 40 mg/kg (without NaOx drinking) for 7 days before harvest.

### Quantitative RT-PCR

Kidney tissue RNA was extracted with Tissue RNA Purification Kit Plus (ES science) according to the manufacturer’s instructions. Hifair 1st Strand cDNA Synthesis SuperMix for qRT-PCR (Yeasen Biotechnology, shanghai) was used for cDNA synthesis. Hieff qPCR SYBR Green Master Mix (Yeasen Biotechnology, shanghai) were used for signal generation detected by CFX96 Touch Real-Time PCR Detection System (Bio-Rad). Relative expression levels were calculated as transcript levels of target genes relative to housekeeping gene GAPDH. The following primers were used:

*il-1β* forward primer ACCTTCCAGGATGAGGACATGA
*il-1β* reverse primer CTAATGGGAACGTCACACACCA
*tnfα* forward primer AAGCCTGTAGCCCACGTCGTA
*tnfα* reverse primer GGCACCACTAGTTGGTTGTCTTTG
*cxcl1* forward primer CTGGGATTCACCTCAAGAACATC
*cxcl1* reverse primer CAGGGTCAAGGCAAGCCTC
*cxcl5* forward primer GGTCCACAGTGCCCTACG
*cxcl5* reverse primer GCGAGTGCATTCCGCTTA
*gapdh* forward primer AGGTCGGTGTGAACGGATTTG
*gapdh* reverse primer TGTAGACCATGTAGTTGAGGTCA

### Statistics

Statistical analysis was performed with Prism 6.0 (GraphPad). Data are presented as mean ± S.E.M. Unpaired Student’s t tests were used to compare two groups. All statistical tests were two tailed, and P values of <0.05 were considered significant.

## Acknowledgments

We thank for the technical support by the Core Facilities, Zhejiang University School of Medicine and the support from Zhejiang Provincial Key Laboratory of Immunity and Inflammatory diseases. In particular, we thank Mrs. Qin Han for their technical assistance on Two-photon Microscopy in the Center of Cryo-Electron Microscopy (CCEM), Zhejiang University; we also thank Mrs. Junli Xuan and Mrs. Shuangshuang Liu for their help in confocal scanning microscopy and image processing.

## REFERENCES

[1] Widmaier, E., H. Raff, and K. Strang, Vander’s Human Physiology (15th Edition). 2019.

[2] Verschuren, E.H.J., C. Castenmiller, D.J.M. Peters, et al. Sensing of tubular flow and renal electrolyte transport. Nat Rev Nephrol, 2020, 16(6): 337–351.

[3] Dantzler, W.H., A.T. Layton, H.E. Layton, et al. Urine-concentrating mechanism in the inner medulla: function of the thin limbs of the loops of Henle. Clin J Am Soc Nephrol, 2014, 9(10): 1781–9.

[4] Sands, J.M. and H.E. Layton. The physiology of urinary concentration: an update. Semin Nephrol, 2009, 29(3): 178–95.

[5] Matlin, K.S. and M.J. Caplan, Seldin and Giebisch’s The Kidney (Fifth Edition). 2013.

[6] Fogazzi, G.B. Crystalluria: a neglected aspect of urinary sediment analysis. Nephrol Dial Transplant, 1996, 11(2): 379–87.

[7] Prescott, L.F. The nephrotoxicity of analgesics. J Pharm Pharmacol, 1966, 18(6): 331–53.

[8] Cisek, L.J. Holding Water: Congenital Anomalies of the Kidney and Urinary Tract, CKD, and the Ongoing Role of Excellence in Plumbing. Adv Chronic Kidney Dis, 2017, 24(6): 357–363.

[9] Wagner, C.A. and N. Mohebbi. Urinary pH and stone formation. J Nephrol, 2010, 23 Suppl 16: S165–9.

[10] Ko, G.J., C.M. Rhee, K. Kalantar-Zadeh, et al. The Effects of High-Protein Diets on Kidney Health and Longevity. J Am Soc Nephrol, 2020, 31(8): 1667–1679.

[11] Dvanajscak, Z., L.N. Cossey, and C.P. Larsen. A practical approach to the pathology of renal intratubular casts. Semin Diagn Pathol, 2020, 37(3): 127–134.

[12] Khan, S.R. and D.J. Kok. Modulators of urinary stone formation. Front Biosci, 2004, 9: 1450–82.

[13] Siener, R. Nutrition and Kidney Stone Disease. Nutrients, 2021, 13(6).

[14] Howles, S.A. and R.V. Thakker. Genetics of kidney stone disease. Nat Rev Urol, 2020, 17(7): 407–421.

[15] Khan, S.R., M.S. Pearle, W.G. Robertson, et al. Kidney stones. Nat Rev Dis Primers, 2016, 2: 16008.

[16] Evan, A.P. Physiopathology and etiology of stone formation in the kidney and the urinary tract. Pediatr Nephrol, 2010, 25(5): 831–41.

[17] Thongprayoon, C., A.E. Krambeck, and A.D. Rule. Determining the true burden of kidney stone disease. Nat Rev Nephrol, 2020, 16(12): 736–746.

[18] Kolter, J., R. Feuerstein, P. Zeis, et al. A Subset of Skin Macrophages Contributes to the Surveillance and Regeneration of Local Nerves. Immunity, 2019, 50(6): 1482–1497 e7.

[19] Whitsett, J.A., S.E. Wert, and T.E. Weaver. Alveolar surfactant homeostasis and the pathogenesis of pulmonary disease. Annu Rev Med, 2010, 61: 105–19.

[20] De Schepper, S., S. Verheijden, J. Aguilera-Lizarraga, et al. Self-Maintaining Gut Macrophages Are Essential for Intestinal Homeostasis. Cell, 2018, 175(2): 400–415 e13.

[21] Hogg, C., A.W. Horne, and E. Greaves. Endometriosis-Associated Macrophages: Origin, Phenotype, and Function. Front Endocrinol (Lausanne), 2020, 11: 7.

[22] Apodaca, G. The uroepithelium: not just a passive barrier. Traffic, 2004, 5(3): 117–28.

[23] Denker, B.M. and E. Sabath. The biology of epithelial cell tight junctions in the kidney. J Am Soc Nephrol, 2011, 22(4): 622–5.

[24] Peerapen, P. and V. Thongboonkerd. Effects of calcium oxalate monohydrate crystals on expression and function of tight junction of renal tubular epithelial cells. Lab Invest, 2011, 91(1): 97–105.

[25] Kawakami, T., J. Lichtnekert, L.J. Thompson, et al. Resident renal mononuclear phagocytes comprise five discrete populations with distinct phenotypes and functions. J Immunol, 2013, 191(6): 3358–72.

[26] Nelson, P.J., A.J. Rees, M.D. Griffin, et al. The renal mononuclear phagocytic system. J Am Soc Nephrol, 2012, 23(2): 194–203.

[27] Zhu, Q., J. He, Y. Cao, et al. Analysis of Mononuclear Phagocytes Disclosed the Establishment Processes of Two Macrophage Subsets in the Adult Murine Kidney. Front Immunol, 2022, 13: 805420.

[28] Regoli, M., E. Bertelli, M. Gulisano, et al. The Multifaceted Personality of Intestinal CX3CR1(+) Macrophages. Trends Immunol, 2017, 38(12): 879–887.

[29] Satpathy, A.T., W. Kc, J.C. Albring, et al. Zbtb46 expression distinguishes classical dendritic cells and their committed progenitors from other immune lineages. J Exp Med, 2012, 209(6): 1135–52.

[30] Rau, W.S. and E. Fromter. Electrical properties of the medullary collecting ducts of the golden hamster kidney. II. The transepithelial resistance. Pflugers Arch, 1974, 351(2): 113–31.

[31] Gonzalez-Mariscal, L., M.C. Namorado, D. Martin, et al. Tight junction proteins ZO-1, ZO-2, and occludin along isolated renal tubules. Kidney Int, 2000, 57(6): 2386–402.

[32] Rescigno, M., M. Urbano, B. Valzasina, et al. Dendritic cells express tight junction proteins and penetrate gut epithelial monolayers to sample bacteria. Nat Immunol, 2001, 2(4): 361–7.

[33] Niess, J.H., S. Brand, X. Gu, et al. CX3CR1-mediated dendritic cell access to the intestinal lumen and bacterial clearance. Science, 2005, 307(5707): 254–8.

[34] Pill, J., B. Kraenzlin, J. Jander, et al. Fluorescein-labeled sinistrin as marker of glomerular filtration rate. Eur J Med Chem, 2005, 40(10): 1056–61.

[35] Barzilai, S., S.K. Yadav, S. Morrell, et al. Leukocytes Breach Endothelial Barriers by Insertion of Nuclear Lobes and Disassembly of Endothelial Actin Filaments. Cell Rep, 2017, 18(3): 685–699.

[36] Paluch, E.K., I.M. Aspalter, and M. Sixt. Focal Adhesion-Independent Cell Migration. Annu Rev Cell Dev Biol, 2016, 32: 469–490.

[37] Mulay, S.R. and H.J. Anders. Crystal nephropathies: mechanisms of crystal-induced kidney injury. Nat Rev Nephrol, 2017, 13(4): 226–240.

[38] Gabarre, P., G. Dumas, T. Dupont, et al. Acute kidney injury in critically ill patients with COVID-19. Intensive Care Med, 2020, 46(7): 1339–1348.

[39] Yu, Y., P. Sikorski, M. Smith, et al. Comprehensive Metaproteomic Analyses of Urine in the Presence and Absence of Neutrophil-Associated Inflammation in the Urinary Tract. Theranostics, 2017, 7(2): 238–252.

[40] Yu, Y., P. Sikorski, C. Bowman-Gholston, et al. Diagnosing inflammation and infection in the urinary system via proteomics. J Transl Med, 2015, 13: 111.

[41] Muller, W.A. Transendothelial migration: unifying principles from the endothelial perspective. Immunol Rev, 2016, 273(1): 61–75.

[42] Carman, C.V. and T.A. Springer. Trans-cellular migration: cell-cell contacts get intimate. Curr Opin Cell Biol, 2008, 20(5): 533–40.

[43] Lin, W.C., K.M. Gowdy, J.H. Madenspacher, et al. Epithelial membrane protein 2 governs transepithelial migration of neutrophils into the airspace. J Clin Invest, 2020, 130(1): 157–170.

[44] Zemans, R.L., S.P. Colgan, and G.P. Downey. Transepithelial migration of neutrophils: mechanisms and implications for acute lung injury. Am J Respir Cell Mol Biol, 2009, 40(5): 519–35.

[45] Azcutia, V., M. Kelm, A.C. Luissint, et al. Neutrophil expressed CD47 regulates CD11b/CD18-dependent neutrophil transepithelial migration in the intestine in vivo. Mucosal Immunol, 2021, 14(2): 331–341.

[46] Gardiner, B.S., D.W. Smith, C.J. Lee, et al. Renal oxygenation: From data to insight. Acta Physiol (Oxf), 2020, 228(4): e13450.

[47] Sands, J.M. and H.E. Layton. Advances in understanding the urine-concentrating mechanism. Annu Rev Physiol, 2014, 76: 387–409.

[48] Brezis, M. and S. Rosen. Hypoxia of the renal medulla--its implications for disease. N Engl J Med, 1995, 332(10): 647–55.

[49] Berry, M.R., R.J. Mathews, J.R. Ferdinand, et al. Renal Sodium Gradient Orchestrates a Dynamic Antibacterial Defense Zone. Cell, 2017, 170(5): 860–874 e19.

