## Supplemental Figures 1-11 for "The specific roles of renal macrophages in monitoring and clearing off intratubular particles"

Fig. S1

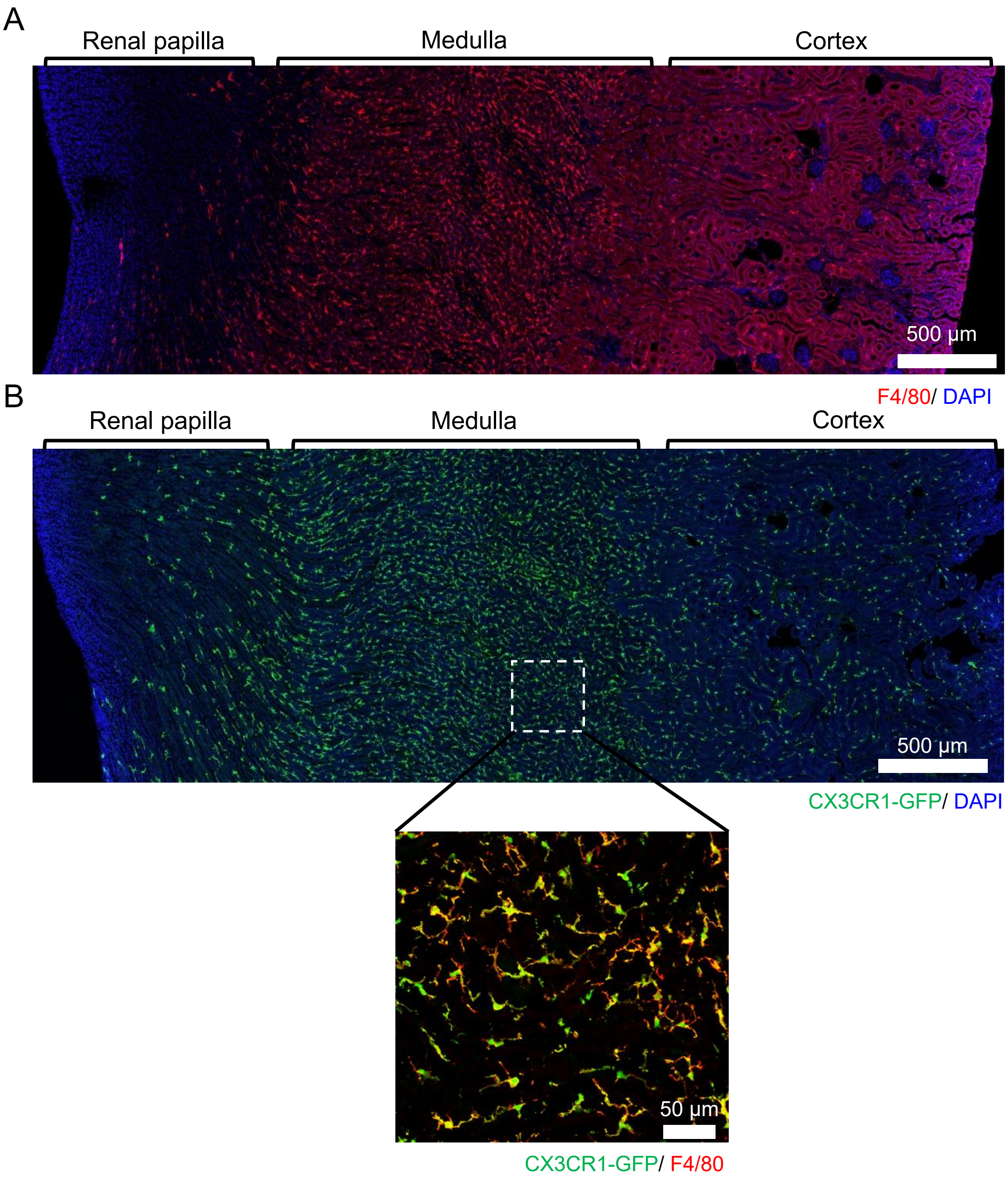


**Figure S1. Anatomical distribution of renal F4/80$^{+}$ and CX3CR1$^{+}$ cells.** (**A**) Immunofluorescent staining for F4/80 in the kidney of adult C57BL/6 mice. (**B**) CX3CR1$^{+}$ cells are shown in the kidney of reporter *Cx3cr1*$^{GFP/+}$ mice. Co-staining for F4/80 shows that almost all CX3CR1$^{+}$ cells in the medulla are F4/80$^{+}$. Representative images from 5 mice of each group.

Fig. S2

A

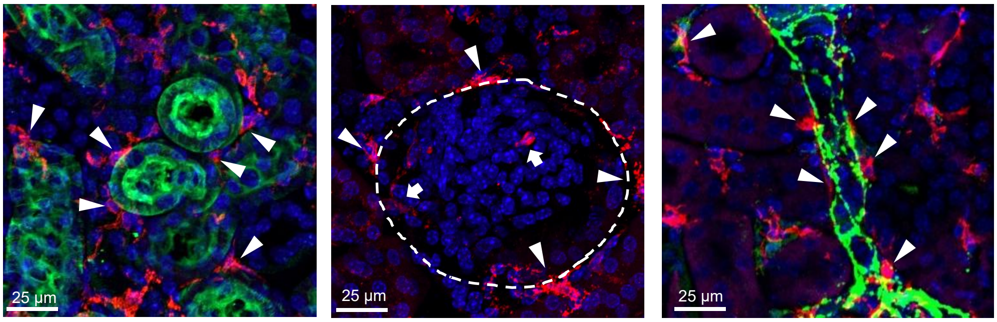


F4/80/ AQP1/ DAPI    F4/80/ DAPI    F4/80/ Tyrosine hydroxylase / DAPI

B

Cortex    Medulla

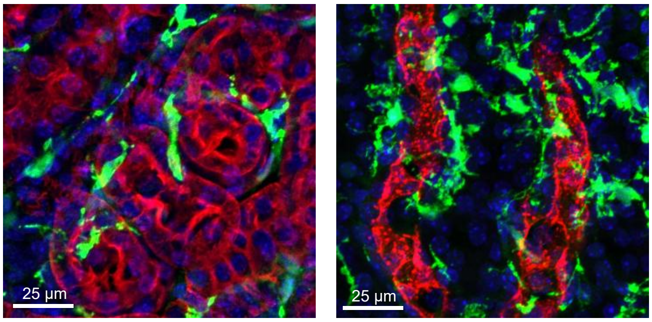


CX3CR1-GFP/ AQP1/ DAPI    CX3CR1-GFP/ AQP2/ DAPI

**Figure S2. Distribution and the morphology of MØ in the microstructures of the kidney.** (**A**) Cortical F4/80⁺ cells were found to be close to AQP1⁺ proximal tubules (arrowhead in the left image), in proximity to the parietal layer of Bowman's capsule (arrowhead in the middle image) and inside glomerulus (arrow in the middle image), and in association with tyrosine hydroxylase-positive sympathetic nerves (arrowhead in the right image). The dashed line delineates the border of a glomerulus. (**B**) Cortical and medullary juxtatubular CX3CR1⁺ cells exhibit morphology distinguishable from each other. AQP1⁺ indicates cortical proximal tubules; AQP2⁺ indicates medullary collecting ducts.

Fig. S3

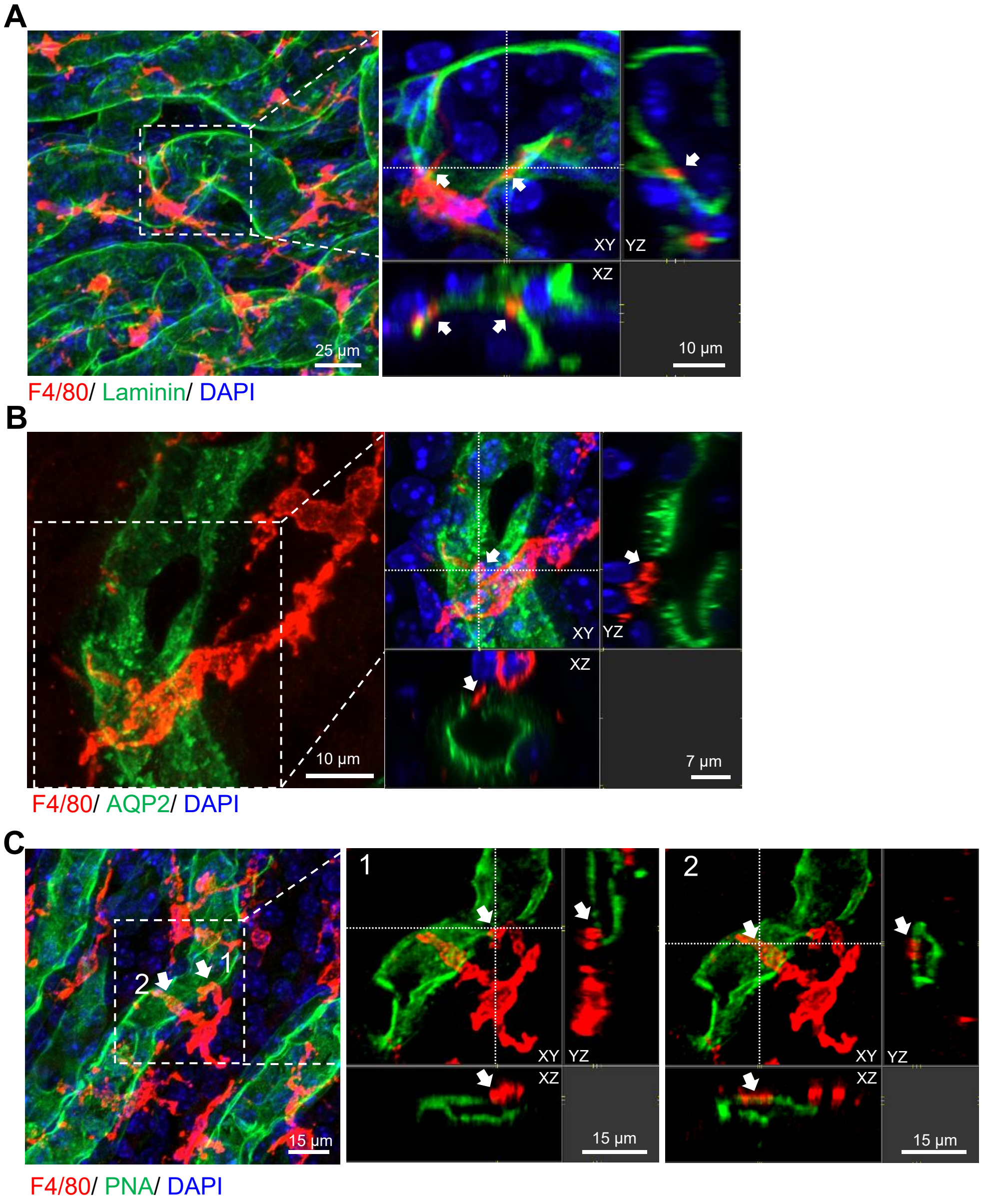


**Figure S3. F4/80⁺ cells extend transepithelial protrusions in the medulla.** (**A**, **B**) Examples of confocal images of medulla sections show that the protrusions of F4/80⁺ cells penetrate laminin⁺ tubular basement membrane (A) and an AQP2⁺ collecting duct (B). (**C**) One juxtatubular F4/80⁺ cell (red) extended two transepithelial protrusions (No. 1 and No. 2) exposed to the PNA⁺ (green) apical surface of a collecting duct. Arrows indicate the parts of protrusions embedded in the epithelium. Z-projections for A, B, and C are 20 µm, 24 µm, and 21 µm, respectively.

Fig. S4

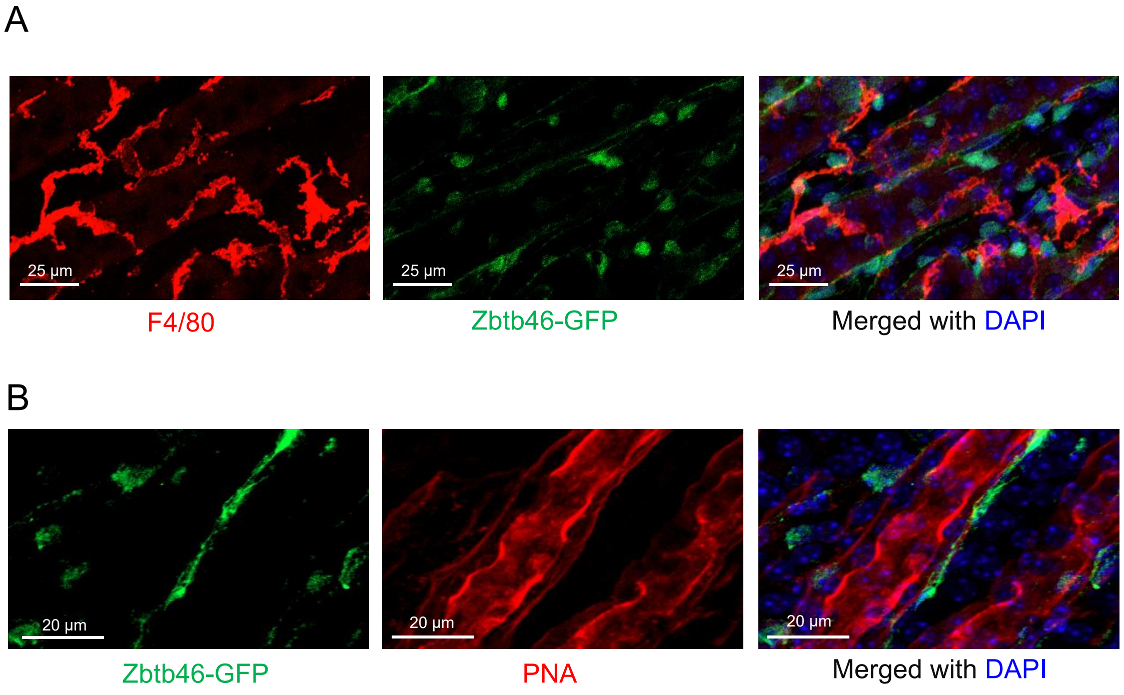


**Figure S4. Medullary DCs do not overlap with F4/80$^{+}$ cells.** (**A**) Immunohistostaining shows that in the renal medulla of *Zbtb46*$^{GFP/+}$ reporter mice, F4/80$^{+}$ cells (red) and classical DCs (green) do not overlap. (**B**) The positional relationship between classical DCs (green) and medullary collecting ducts (red). Representative pictures from 3 mice.

Fig. S5

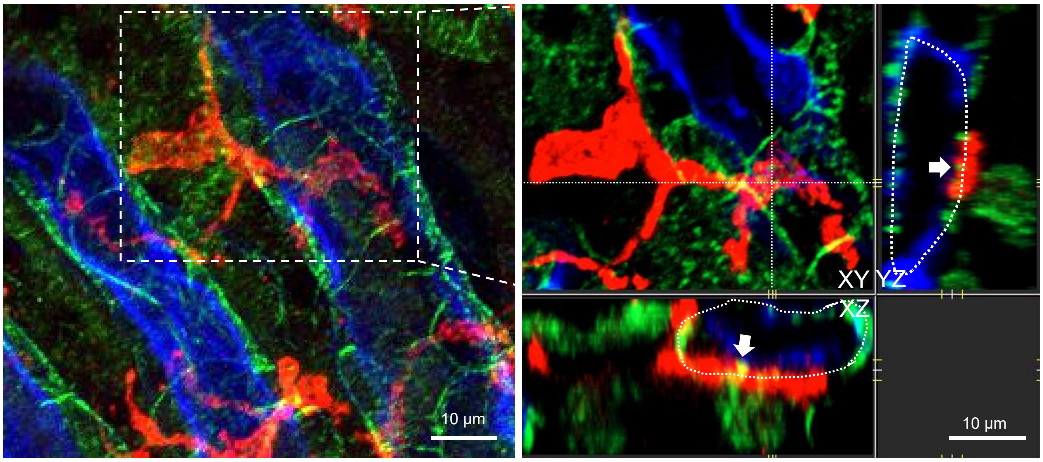


F4/80/ β-catenin/ PNA

**Figure S5. MØ make transepithelial protrusions through a transcellular route.** Representative confocal image of a transepithelial protrusion derived from a MØ (red) with epithelial border protein β-catenin (green) on epithelium (blue). The arrows indicate intact border structures (green) wrapped around by the transepithelial protrusion. Z-projections of 22 µm.

Fig. S6

A

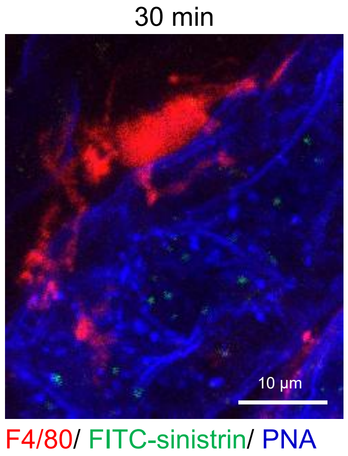


B

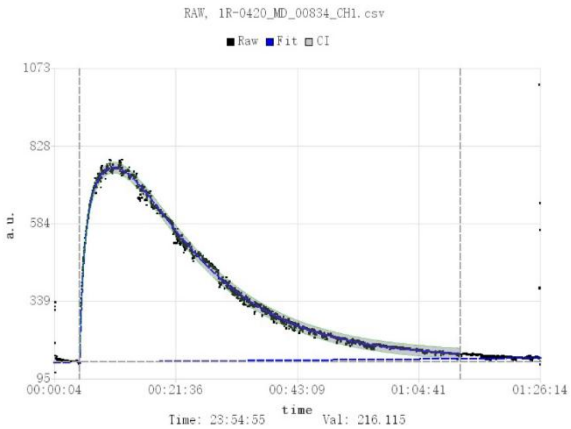


**Figure S6. Kinetics of FITC-sinistrin removal after i.v. administration.** (**A**) The medullary collecting ducts (blue) were examined 30 min post i.v. infusion of FITC-sinistrin. Z-projections of 21 µm. Note that at this time point, FITC-sinistrin (green) had showed up in the lumen of collecting ducts, but had not been uptaken by MØ yet. (**B**) A typical clearance curve (Time vs. Florescence intensity) of FITC-sinistrin in naïve mouse, as measured by a miniaturized fluorescent densitometer (MediBeacon, Germany) attached on shaved skin of the back.

Fig. S7

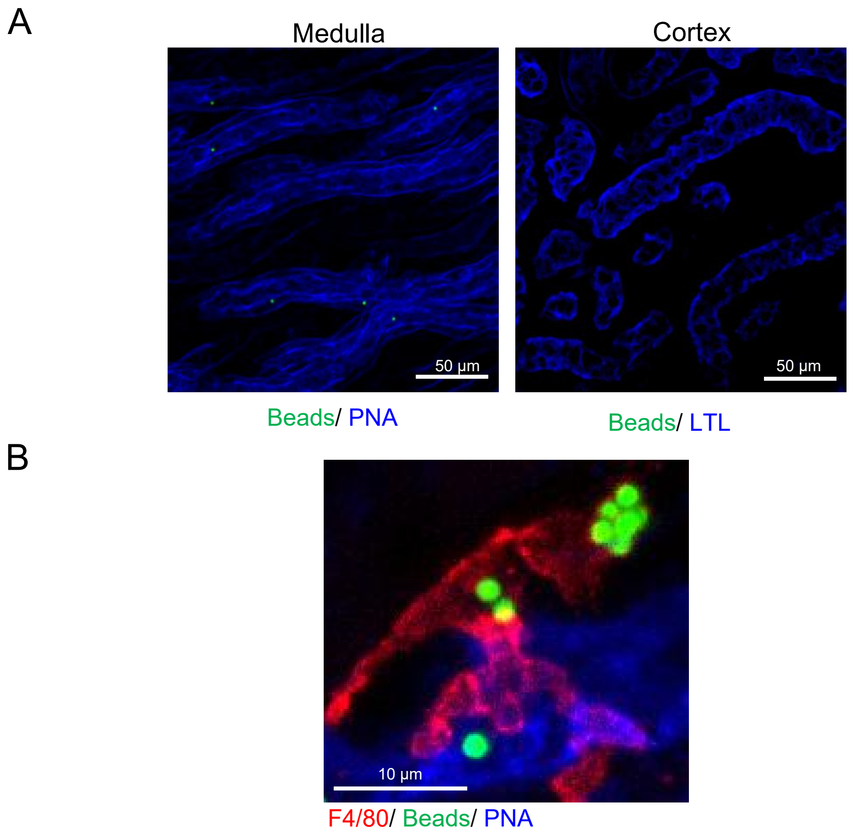


**Figure S7. The distribution of latex beads after intrapelvic administration.** (**A**) Two hours after intrapelvic injection, fluorescent beads localized in the PNA$^{+}$ collecting ducts but not lotus tetragonolobus lectin (LTL)-positive proximal tubules. (**B**) Confocal images show that 12 hr after intrapelvic injection, a MØ (red) in the proximity of a medullary collecting duct (blue) had uptaken multiple beads (green) which had been transported to the soma outside of the tubule. Representative pictures from at least 6 mice.

Fig. S8

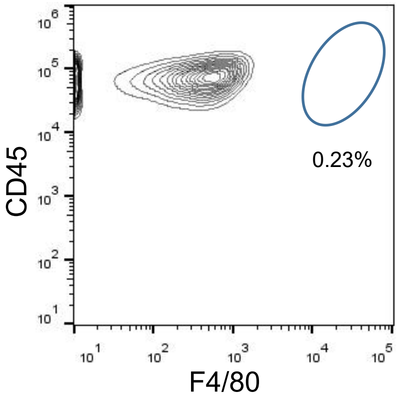


**Figure S8. F4/80hi cells are absent in the blood.** After lysis of erythrocytes from the blood, leukocytes were analyzed by flow cytometry. Representative contour plot of 3 mice.

Fig S9

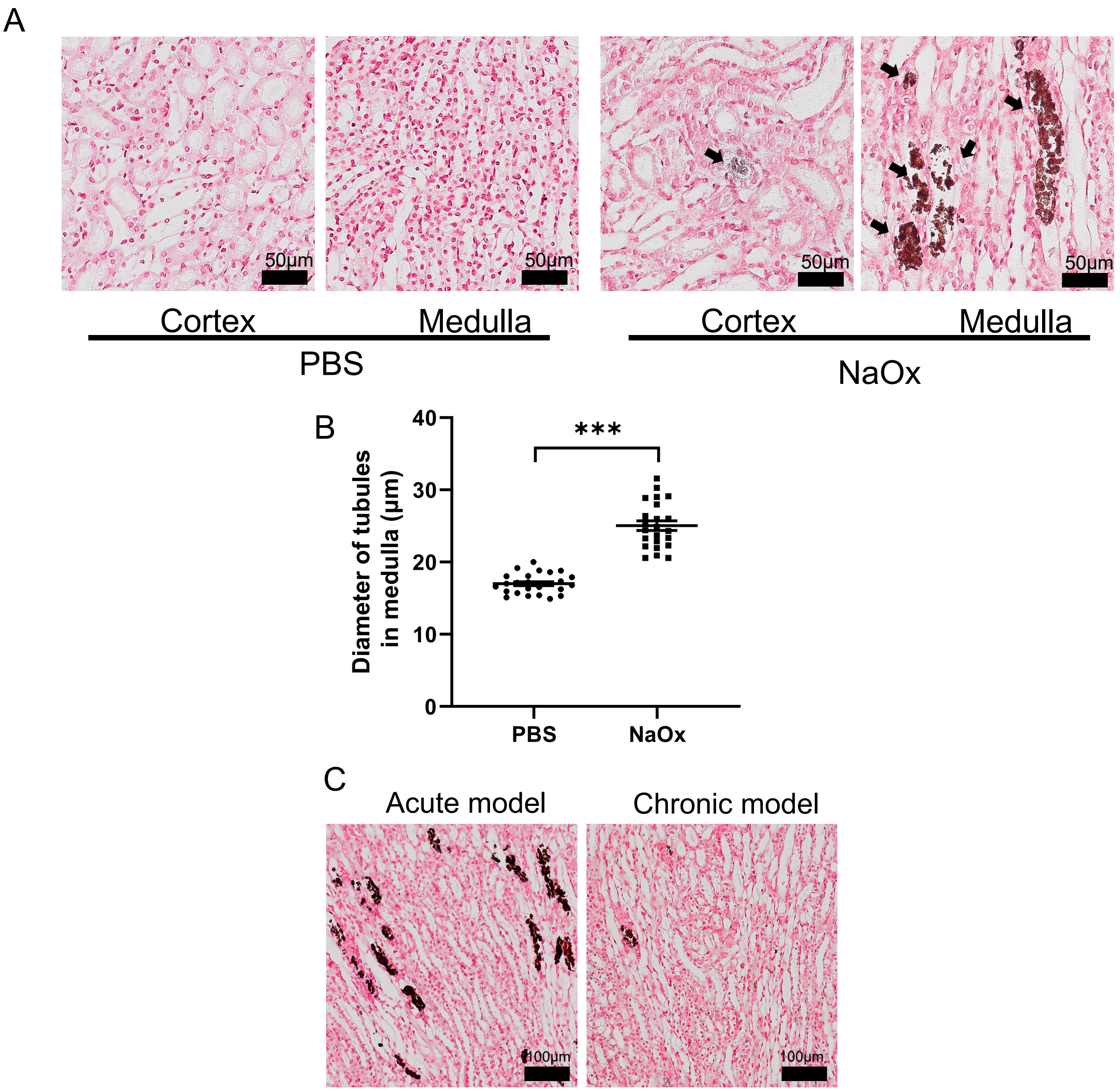


**Figure S9. Formation of kidney stones in two nephrolithiasis models.** (**A**, **B**) An acute nephrolithiasis model by i.p. injection of sodium oxalate (NaOx) (100 mg/kg) plus 3% NaOx in drinking water to C57BL/6 mice. The mice were examined 24 hr later. The controls were i.p. treated with PBS and drank normal water. (A) Representative Von Kossa staining shows the CaOx crystal deposition in the cortex and medulla. (B) The diameters of medullary tubules were measured. Each dot indicates an average from a 290 μm×290 μm FOV. n = 4. ***$P<0.005$ by two-tailed unpaired t test. Data are depicted as mean ± SEM. (**C**) Representative Von Kossa staining of renal medulla of C57BL/6 mice induced with an acute model as above or a chronic model of nephrolithiasis by being i.p. treated with NaOx (40 mg/kg) for 7 days.

Fig. S10

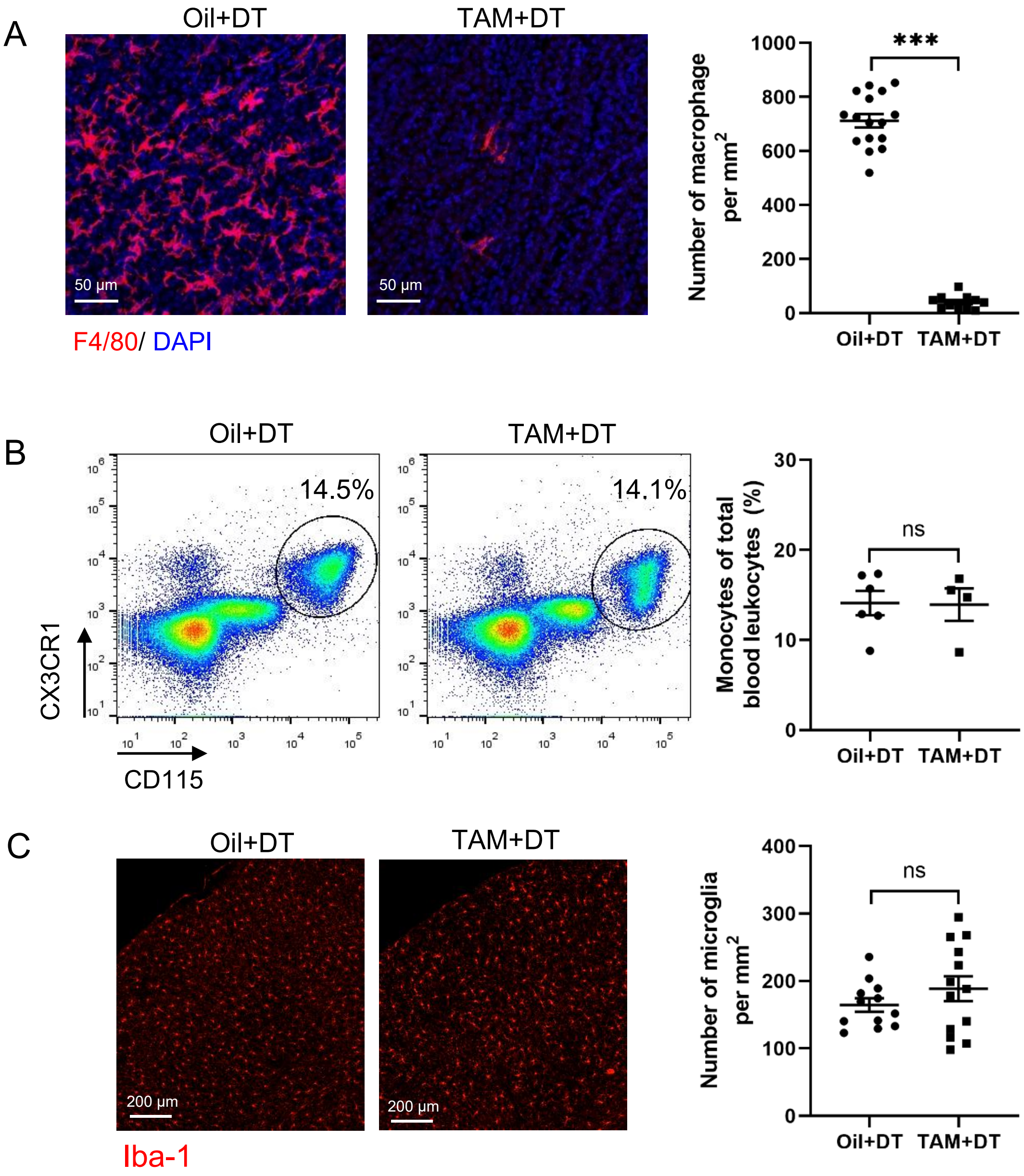


**Figure S10. Selective depletion of MØ in the renal medulla**. Female *Cx3cr1$^{CreER/+}$:iDTR* mice were first treated with corn oil or tamoxifen. After that, 2 shots of intravesical DT was administered 24 hr apart. The following parameters were measured 2 days after the 2$^{nd}$ DT application: (**A**) the density of medullary MØ, evaluated by immunohistostaining. Each dot in the quantification panel indicates the sum of MØ from a 319 µm×319 µm FOV. n = 3; (**B**) the abundance of blood monocytes evaluated by flow cytometry analysis and; (**C**) the density of microglia in brain cortex, as evaluated by immunohistostaining of Iba-1. Each dot in the quantification panel indicates the sum of microglia from a 1280 µm×1280 µm FOV. n = 3. Data are depicted as mean ± SEM. ***$P$ < 0.005 by two-tailed unpaired t test. ns, not significant.

Fig. S11

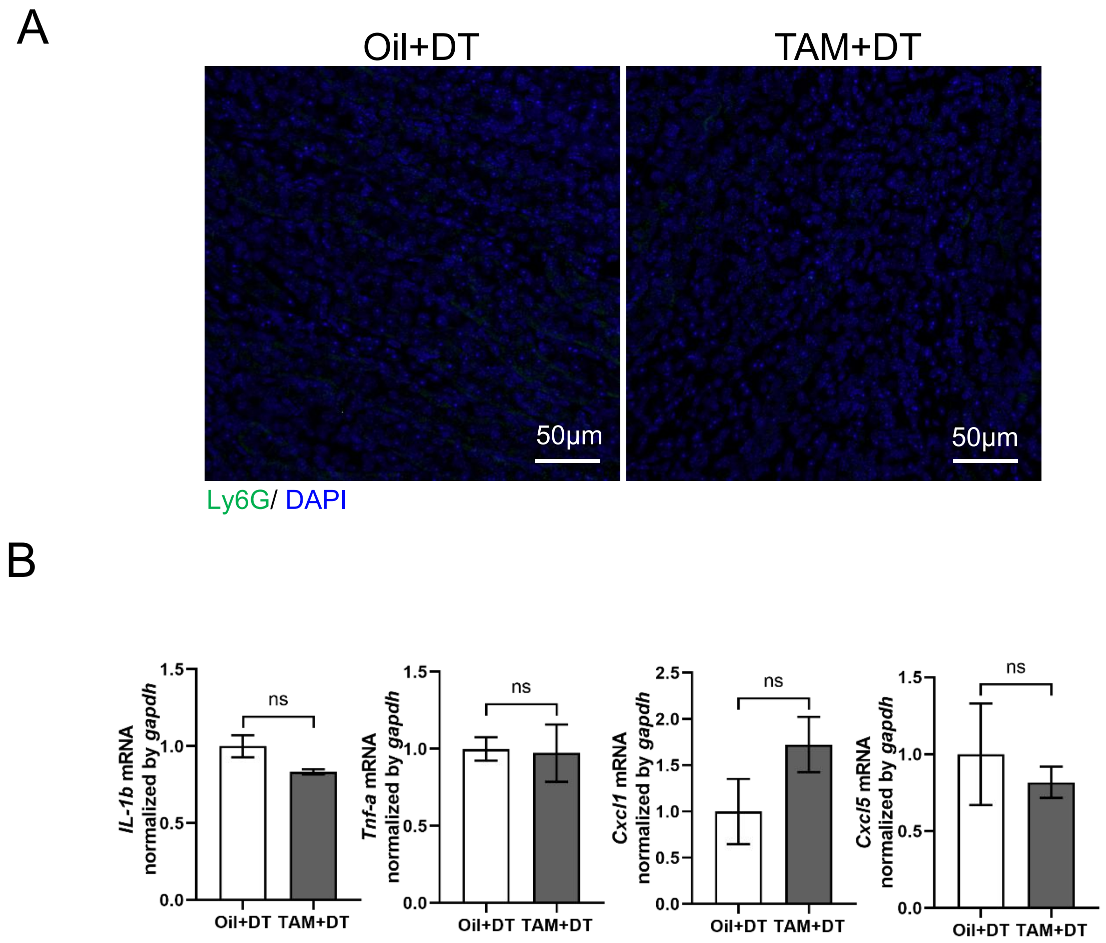


**Figure S11. MØ depletion itself did not induce overt inflammation**. *Cx3cr1*$^{CreER/+}$:*iDTR* mice were first treated with corn oil or tamoxifen; after that, intravesical application of DT was performed, as depicted in Fig. 4D. Two days after DT application, (**A**) immunohistostaining images for neutrophils in the medulla show no infiltration of Ly6G$^+$ cells (green) in either group (representative of 4 mice in each group); (**B**) relative expression of the indicated inflammatory cytokines/chemokines in the kidneys was assessed by quantitative RT-PCR. n = 7. ns, not significant by two-tailed unpaired t test.
